# Orthohantavirus-related Proteases as Therapeutic Targets: Opportunities for Antiviral Drug Development

**DOI:** 10.64898/2026.05.12.724423

**Authors:** Jakub M. Tomczak, Ewelina Węglarz-Tomczak

## Abstract

Orthohantaviruses cause severe human diseases including hemorrhagic fever with renal syndrome (HFRS) and hantavirus cardiopulmonary syndrome (HCPS), with case fatality rates up to 40%. No FDA-approved therapeutics are currently available, highlighting urgent need for drug development following recent outbreak events. We systematically examined host protease dependencies in hantavirus replication, focusing on Signal Peptidase (SP) and Signal Peptide Peptidase (SPP) essential for viral glycoprotein maturation. Through comprehensive database mining and molecular docking analysis, we identified six potential protease inhibitors, with Compound E achieving the highest binding confidence score (−0.28) against SPP. Our analysis reveals that targeting host ER proteases represents a viable antiviral strategy, providing a systematic framework for protease-targeted antihantavirus drug development and identifying specific lead compounds for experimental validation.

## 1. Introduction

Orthohantaviruses (hereafter referred to as hantaviruses), in the family Hantaviridae, are human pathogenic viruses causing severe diseases with high mortality rate and represent therefore a serious challenge for public health (Afzal et al., 2023; Hart & Bennett, 1999). These enveloped, negative-sense RNA viruses belong to the order Bunyavirales and are categorized into two major groups based on their geographical distribution and associated disease pathology. Their genomes consist of three negative-sense single-stranded RNA segments designated small (S), medium (M), and large (L), encoding the nucleocapsid protein (N), glycoprotein precursor (GPC), and RNA-dependent RNA polymerase (RdRp), respectively. Human infection primarily occurs through inhalation of aerosolized excreta from infected rodent reservoirs. Old World hantaviruses, including Hantaan virus (HTNV) and Seoul virus (SEOV), predominantly cause hemorrhagic fever with renal syndrome (HFRS), whereas New World hantaviruses such as Andes virus (ANDV) and Sin Nombre virus (SNV) cause hantavirus cardiopulmonary syndrome (HCPS), with up to 40% mortality (Khaiboullina et al., 2005).

Proteases are biological targets playing pivotal roles in multiple diseases (Turk, 2006), such as, cancer (Weglarz-Tomczak et al., 2018), neurodegenerative diseases (Senkowska & Weglarz-Tomczak, 2026), and viral infections (Agbowuro et al., 2018). However, unlike coronaviruses (Weglarz-Tomczak et al., 2021), flaviviruses (Teramoto et al., 2023), or retroviruses (Dunn et al., 2002), orthohantaviruses do not encode a dedicated viral protease for polyprotein processing. As a result, antiviral development must focus on alternative enzymatic or proteolytic vulnerabilities, particularly host proteases involved in viral maturation and entry.

## 2. The Mechanism of Hantavirus Cell Infection

A central feature of hantavirus biology and pathogenesis is the synthesis and processing of the viral glycoprotein precursor (GPC), which gives rise to the mature envelope glycoproteins Gn and Gc. These glycoproteins are essential for viral attachment, entry, assembly, and immune evasion (Acuna et al., 2014; Cifuentes-Munoz et al., 2014; Lober et al., 2001; Shi et al., 2016).

The hantavirus M genomic segment encodes a single glycoprotein precursor polyprotein. Following translation in the rough endoplasmic reticulum (ER), this precursor undergoes proteolytic processing mediated primarily by host-cell proteases rather than viral enzymes. The most important enzyme involved is the cellular signal peptidase (SP), an ER-associated protease complex responsible for cotranslational cleavage of membrane and secretory proteins. SP cleaves the glycoprotein precursor to generate the mature Gn and Gc glycoproteins (Lober et al., 2001; Meier et al., 2021; Muyangwa et al., 2015), as illustrated in Figure 1(a).

**Figure 1.**
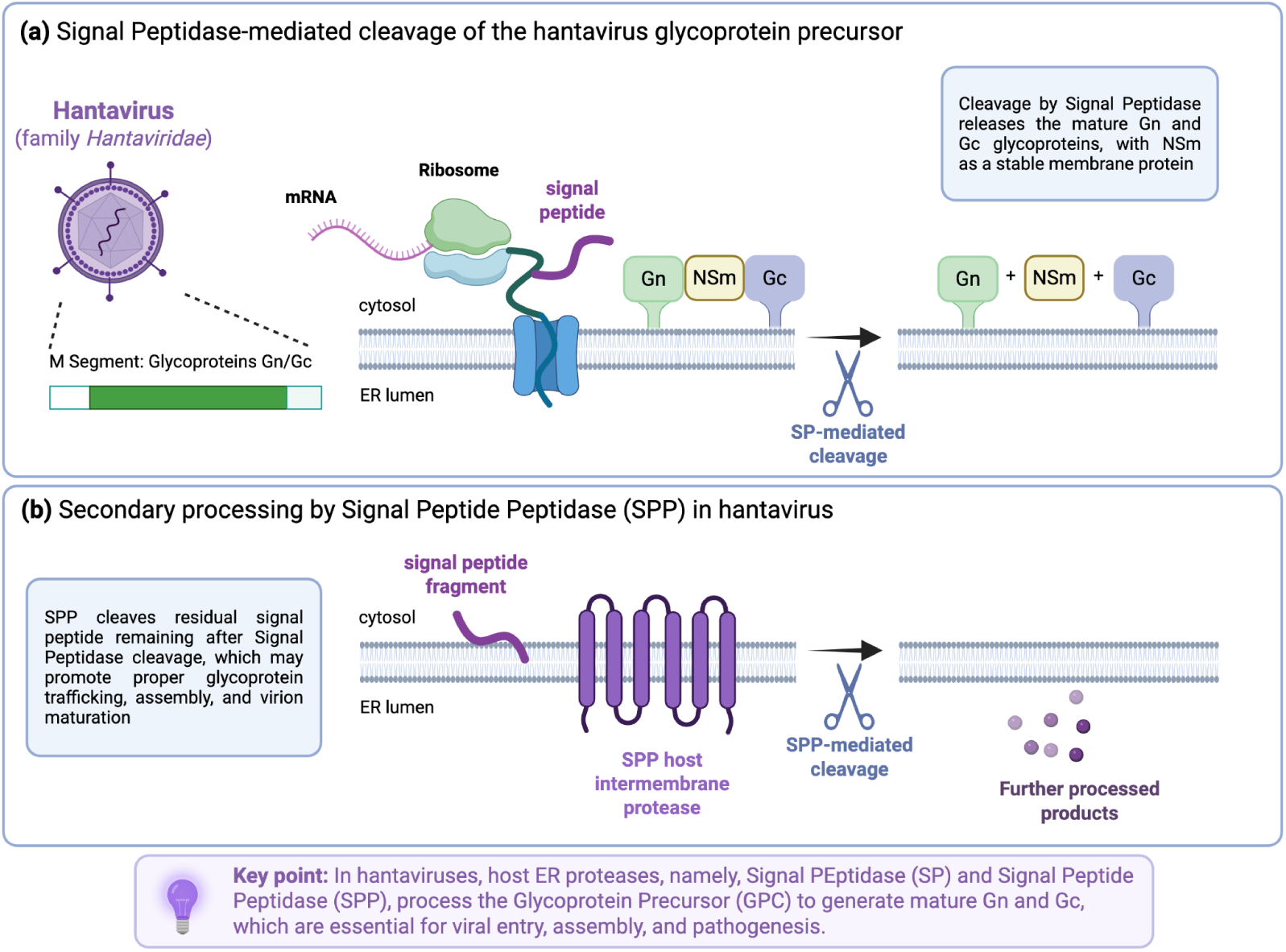
Proteolytic processing of the hantavirus glycoprotein precursor in hantavirus infection. Hantaviruses use host endoplasmic reticulum (ER) proteases to process the glycoprotein precursor (GPC) encoded by the viral M segment into mature Gn and Gc glycoproteins. (a) Cotranslational processing of the hantavirus GPC by the host signal peptidase complex (SP) during translocation into the ER membrane. The M segment encodes a single glycoprotein precursor polyprotein that contains the Gn and Gc envelope glycoproteins separated by a nonstructural membrane-associated protein (NSm). Following ribosome-mediated translation and insertion into the ER membrane, signal peptidase cleaves the precursor at specific signal peptide cleavage sites, generating the mature membrane-associated Gn and Gc glycoproteins required for virion assembly, receptor interaction, and membrane fusion. (b) Secondary processing mediated by signal peptide peptidase (SPP), an intramembrane aspartyl protease localized within the ER membrane. Following SP cleavage, residual signal peptide fragments remaining within the membrane are further processed by SPP, generating smaller peptide products and potentially contributing to optimal glycoprotein maturation, intracellular trafficking, and virion morphogenesis. Together, these host ER proteases constitute essential components of the hantavirus replication cycle and represent potential targets for antiviral intervention. Created with BioRender-style scientific illustration elements. *Created in* https://BioRender.com

In several bunyaviruses, including hantaviruses and related orthobunyaviruses, signal peptide peptidase (SPP), an intramembrane aspartyl protease, also contributes to glycoprotein maturation (Schwake et al., 2022). As shown in Figure 1(b), SPP processes residual signal peptide fragments remaining after SP cleavage and may influence proper glycoprotein trafficking, assembly, and virion maturation. Thus, the maturation pathway of hantaviral glycoproteins depends heavily on host ER proteolytic machinery.

The mature Gn and Gc glycoproteins subsequently form heterodimeric complexes that localize predominantly to the Golgi apparatus, where virion assembly occurs. Gn is primarily involved in receptor interaction and shielding of the fusion machinery, whereas Gc functions as the viral fusion protein that mediates membrane fusion during host-cell entry. These glycoproteins facilitate viral attachment to endothelial cells through interactions with β-integrins and other host receptors, contributing to the characteristic vascular dysfunction observed in hantavirus disease (Afzal et al., 2023).

Endothelial infection and dysregulation are central to hantavirus pathogenesis. Although hantaviruses are generally non-cytopathic, infection induces alterations in endothelial barrier function, leading to increased vascular permeability. This vascular leakage underlies the major clinical manifestations of hantavirus disease, including hantavirus pulmonary syndrome (HPS) and hemorrhagic fever with renal syndrome (HFRS) (Hart & Bennett, 1999). Immune-mediated mechanisms, including excessive cytokine production, activation of cytotoxic T lymphocytes, and complement activation, further amplify endothelial dysfunction and tissue injury .

Importantly, the proteolytic processing of the glycoprotein precursor is performed by host proteases and not by the mature Gn glycoprotein itself (Acuna et al., 2014; Cifuentes-Munoz et al., 2014). Gn functions primarily as a structural and attachment glycoprotein rather than as a protease enzyme. Therefore, host-cell proteolytic systems are indispensable for the generation of infectious hantavirus particles and represent potential targets for antiviral intervention.

Collectively, these observations highlight host proteases as potentially important therapeutic targets in hantavirus infection. In particular, signal peptidase (SP) and signal peptide peptidase (SPP) are of considerable interest because they are directly involved in the proteolytic maturation of the hantavirus glycoprotein precursor (GPC), generating the mature Gn and Gc glycoproteins as well as additional processed peptide products required for efficient viral assembly and maturation. Disruption of these host ER-associated proteolytic pathways could therefore impair the production of infectious virions and limit viral propagation.

Additional classes of host proteases may also contribute to hantavirus entry and intracellular trafficking. Among these, endosomal and lysosomal proteases, particularly cathepsins, have attracted attention due to their established roles in the entry mechanisms of several enveloped viruses (Brix, 2018). Cathepsin-mediated processing has been proposed to facilitate membrane fusion or endosomal escape in certain hantavirus species. However, current evidence does not support a universal dependence of hantaviruses on cathepsin activity, and the available data remain limited and, in some cases, virus-specific. Consequently, while endosomal and lysosomal proteases remain potentially relevant antiviral targets, here, we do not further examine this protease class in detail.

## 3. Proteolytic Processing Pathways Relevant to Orthohantavirus

### 3.1 Signal Peptidase (SP)

Signal peptidase (SP), more precisely the signal peptidase complex (SPC), is a membrane-associated host protease located in the endoplasmic reticulum (ER) (Auclair et al., 2012; Paetzel et al., 2002; Zanotti et al., 2022). Its primary function is cleavage of N-terminal signal peptides from nascent secretory and membrane proteins during cotranslational translocation into the ER lumen.

In orthohantaviruses, SP is critically involved in processing of the viral GPC, encoded by the M segment. The precursor is cleaved at a highly conserved pentapeptide motif, typically WAASA, generating the mature envelope glycoproteins: Gn (formerly G1), and Gc (formerly G2). This cleavage event is essential for: (i) proper glycoprotein folding, (ii) oligomerization, (iii) Golgi trafficking, (iv) virion assembly, (v) infectivity. Failure of SP-mediated processing results in noninfectious viral particles or impaired glycoprotein transport.

Signal peptidase belongs to the class of serine proteases, more specifically, ER-type signal peptidases (Enzyme Commission: EC 3.4.21.89). Unlike classical trypsin-like serine proteases, the catalytic mechanism of eukaryotic SP relies on a Ser–Lys catalytic dyad rather than the canonical Ser–His–Asp triad. Catalytic residues are located primarily within the SEC11 subunit of the complex.

The mammalian signal peptidase complex is a hetero-oligomeric membrane complex composed of multiple subunits: SEC11A / SEC11C (catalytic subunit), SPCS1 (structural/support), SPCS2 (substrate interaction), SPCS3 (complex stabilization). The catalytic center is embedded within the ER membrane interface, allowing recognition of signal peptides as they emerge from the Sec61 translocon. Recent electron microscopy studies have substantially improved understanding of SPC architectures (Liaci et al., 2021). The 3D structures of SPCs are presented in Figure 2.

**Figure 2.**
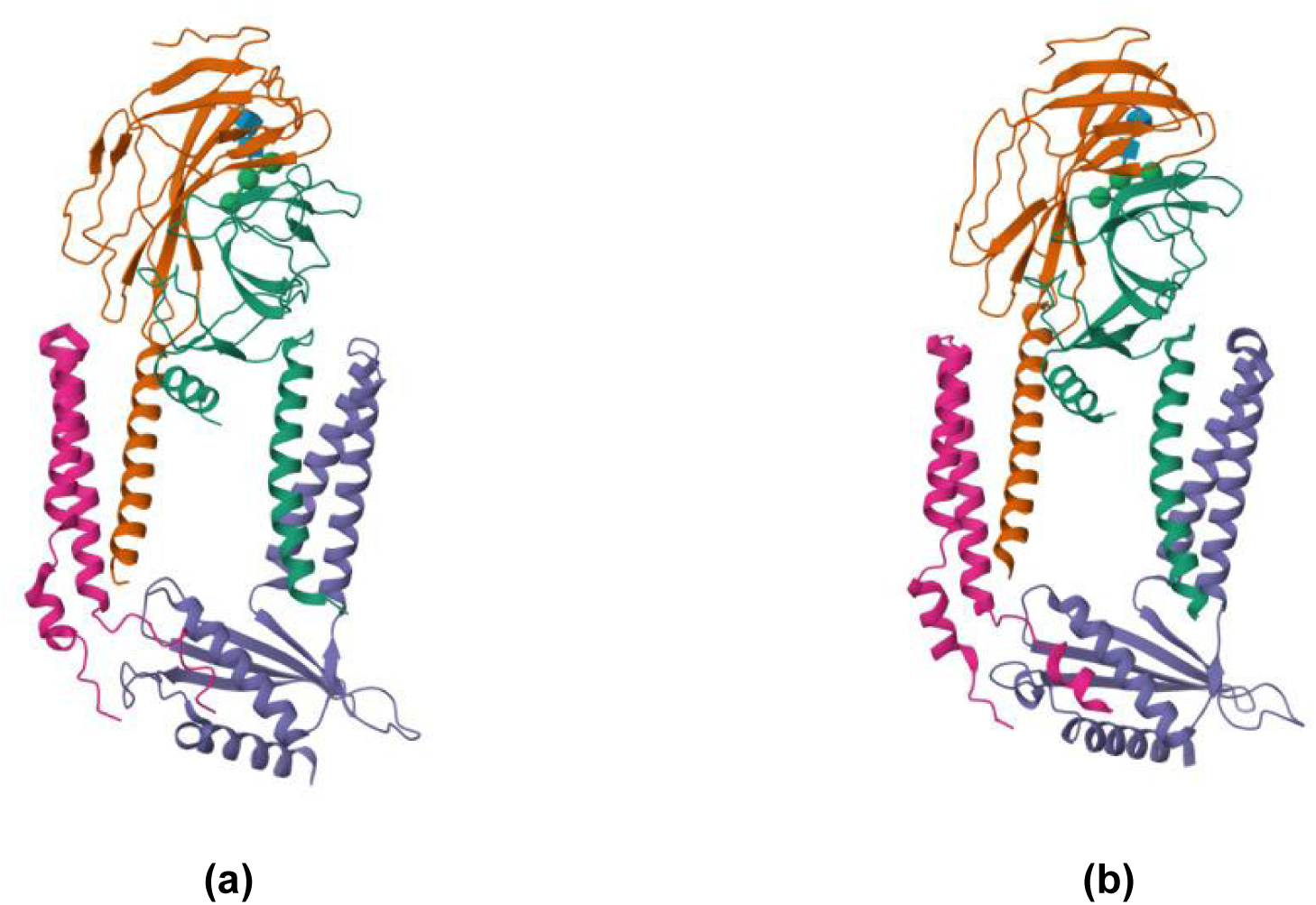
Structural organization of human Signal Peptidase Complex paralogs involved in endoplasmic reticulum protein processing. (a) Human Signal Peptidase Complex Paralog A (SPC-A), PDB ID: 7P2P; (b) Human Signal Peptidase Complex Paralog C (SPC-C), PDB ID: 7P2Q. The signal peptidase complex (SPC) is an endoplasmic reticulum (ER)-resident membrane-associated protease complex responsible for cotranslational cleavage of signal peptides from nascent secretory and membrane proteins. SPC-A and SPC-C represent two distinct human paralogous complexes that differ in subunit composition while maintaining a conserved catalytic architecture. In both structures, the complexes are composed of multiple transmembrane helices supporting luminal catalytic domains responsible for signal peptide recognition and cleavage. The catalytic SEC11 subunit, which contains the serine-lysine catalytic dyad essential for proteolytic activity, is associated with accessory SPC subunits that stabilize the complex and contribute to substrate specificity and membrane organization.

The hantavirus M segment encodes a single glycoprotein precursor translated into the ER membrane. Host SPC cleaves the precursor co-translationally at the conserved WAASA motif.

The cleavage process is required for generation of: mature Gn/Gc heterodimers, proper virion envelope assembly, surface lattice formation. Because Gn/Gc are directly involved in receptor binding and membrane fusion, SPC indirectly controls viral infectivity. Experimental mutagenesis of the WAASA motif disrupts cleavage and abolishes production of infectious particles (Lober et al., 2001).

### 3.2 Signal Peptide Peptidase (SPP)

Signal peptide peptidase (SPP) is an intramembrane-cleaving protease located in the ER membrane (Lemberg et al., 2002; Voss et al. 2013; Weihofen et al., 2002). Following cleavage of signal peptides by SPC, residual peptide fragments embedded within the membrane are further processed by SPP. SPP therefore acts downstream of signal peptidase in protein maturation pathways. Unlike SPC, which cleaves luminally exposed peptide bonds, SPP catalyzes intramembrane proteolysis within hydrophobic transmembrane domains.

SPP belongs to the family of aspartyl intramembrane proteases (Enzyme Commission: EC 3.4.23.B24). SPP is mechanistically related to presenilins, and γ-secretase family proteases. SPP’s catalysis depends on two conserved transmembrane aspartate residues.

SPP is a multi-pass transmembrane ER protein typically containing 7-9 transmembrane helices, catalytic YD and GXGD motifs, and intramembrane active site cavity. The active site is embedded within the lipid bilayer, allowing hydrolysis of peptide bonds inside transmembrane helices. Key conserved motifs include the YD motif and GXGD motif. These motifs are hallmarks of intramembrane aspartyl proteases.

Recent advances in cryo-electron microscopy have enabled structural characterization of human members of the SPP/SPPL family. Particularly important is the cryo-EM structure of human signal peptide peptidase-like 2A (SPPL2A), which provides direct insight into the architecture of intramembrane aspartyl proteases belonging to the SPP family (Huag et al., 2025), see Figure 3.

**Figure 3.**
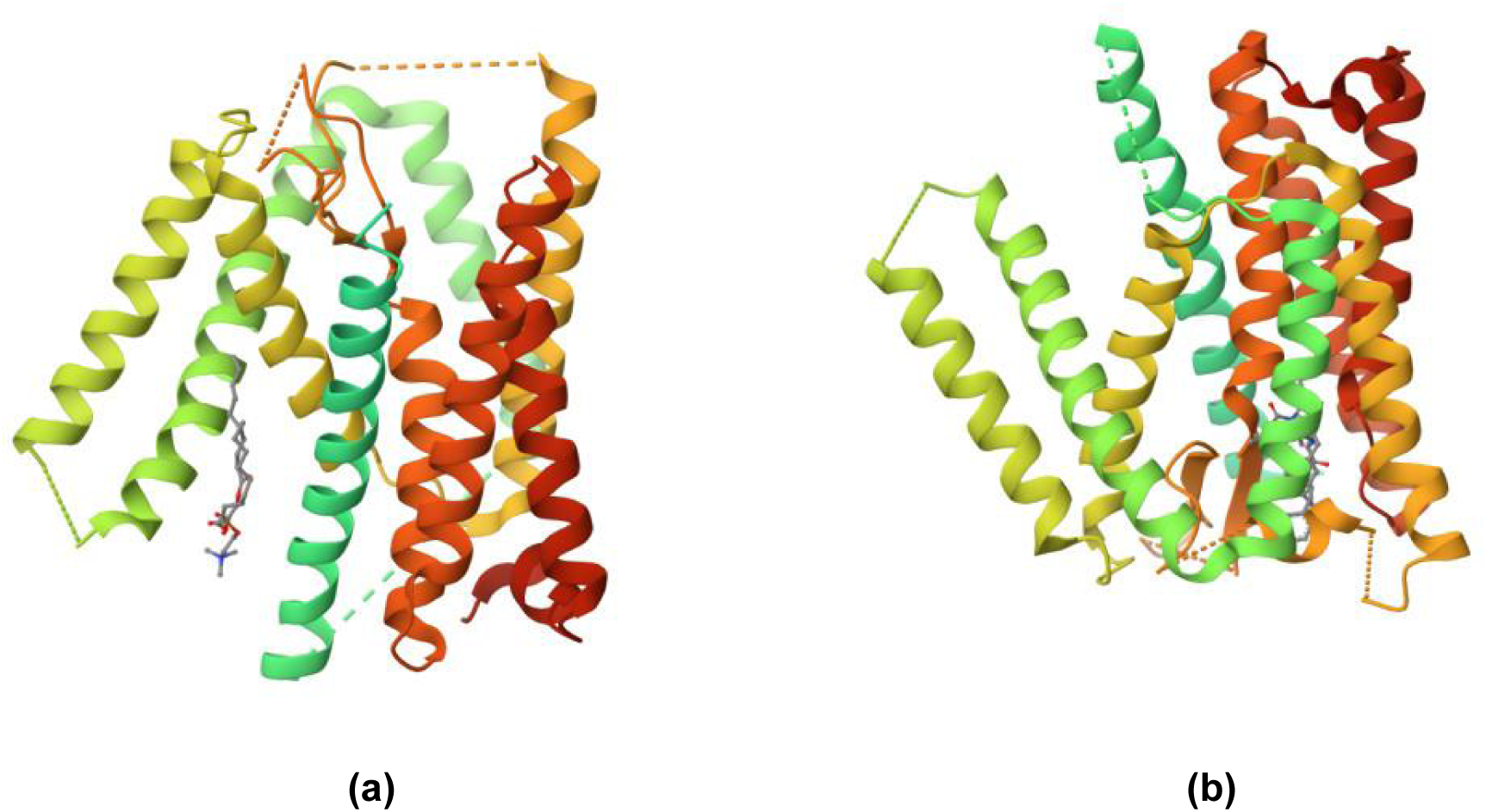
Structural organization of human representative structures of SPP-family intramembrane aspartyl proteases. (a) Human signal peptide peptidase like 2A (SPPL2a) in complex with 3-SN-PHOSPHATIDYLCHOLINE, PDB ID: 9K92; (b) Human signal peptide peptidase like 2A (SPPL2a) in complex with L-685,458, PDB ID: 9K93. SPPL2A belongs to the signal peptide peptidase (SPP)/presenilin family of intramembrane-cleaving aspartyl proteases and exhibits the characteristic multi-pass transmembrane helical architecture associated with this enzyme class. The structures reveal the membrane-embedded catalytic cavity formed by conserved transmembrane helices containing the catalytic YD and GXGD motifs, which coordinate intramembrane proteolysis within the lipid bilayer. Comparison between the apo-like lipid-associated state (9K92) and inhibitor-bound conformation (9K93) illustrates substrate-access pathways and inhibitor engagement within the hydrophobic catalytic pocket. These structural features provide mechanistic insight into SPP-family catalysis and support structure-guided development of inhibitors targeting intramembrane aspartyl proteases potentially involved in viral glycoprotein processing and ER-associated host-virus interactions.

The structure demonstrates a multi-pass transmembrane organization, a formation of a membrane-embedded catalytic cavity, positioning of the conserved catalytic motifs, and substrate-access pathways within the lipid bilayer. These findings significantly improve mechanistic understanding of intramembrane proteolysis and provide a more accurate framework for inhibitor development targeting SPP-family enzymes.

The precise role of SPP in hantavirus replication remains incompletely characterized. However, several mechanistic possibilities make SPP biologically relevant:

● *Processing of residual viral signal peptides*: After SPC-mediated cleavage of hantavirus GPC, remaining membrane-embedded signal peptides may require SPP processing.
● *Regulation of ER-associated degradation (ERAD)*: SPP participates in membrane protein turnover and ER quality control, pathways potentially exploited during viral infection.
● *Immune modulation*: SPP contributes to generation of certain antigenic peptides for MHC presentation, potentially influencing antiviral immunity.
● *Control of membrane proteostasis*: Hantavirus glycoproteins heavily remodel ER and Golgi membranes, therefore, SPP-associated quality-control pathways may influence viral maturation efficiency.

Although direct experimental evidence specifically linking SPP inhibition to reduced hantavirus replication remains limited, the enzyme represents a plausible host dependency factor deserving further study.

### 3.3 Comparison Between SPC and SPP

Although both SPC and SPP participate in sequential stages of membrane protein processing within the endoplasmic reticulum, their mechanistic roles and therapeutic relevance in orthohantavirus infection differ substantially. SPC directly mediates cleavage of the hantavirus glycoprotein precursor at the conserved WAASA motif, thereby generating the mature Gn and Gc envelope glycoproteins required for viral assembly and infectivity. Consequently, SPC represents a proximal and mechanistically validated host dependency factor in the hantavirus life cycle. In contrast, SPP functions downstream of SPC by catalyzing intramembrane cleavage of residual signal peptide fragments. While its precise contribution to hantavirus replication remains incompletely understood, SPP may indirectly influence viral maturation through modulation of ER proteostasis, membrane protein turnover, and antigen presentation pathways. Thus, SPP currently represents a more exploratory antiviral target.

From a structural and biochemical perspective, the two enzymes belong to fundamentally different protease classes. SPC is a membrane-associated serine protease utilizing a Ser–Lys catalytic dyad, whereas SPP is an intramembrane aspartyl protease containing conserved catalytic aspartates within transmembrane helices. These mechanistic differences may have important implications for inhibitor design, selectivity, and toxicity profiles.

Overall, SPC appears to constitute the more biologically validated target in orthhantavirus infection, whereas SPP remains of interest primarily for its broader role in ER-associated membrane processing and host–virus interactions. The comparison between the two is presented in Table 1.

**Table 1.**
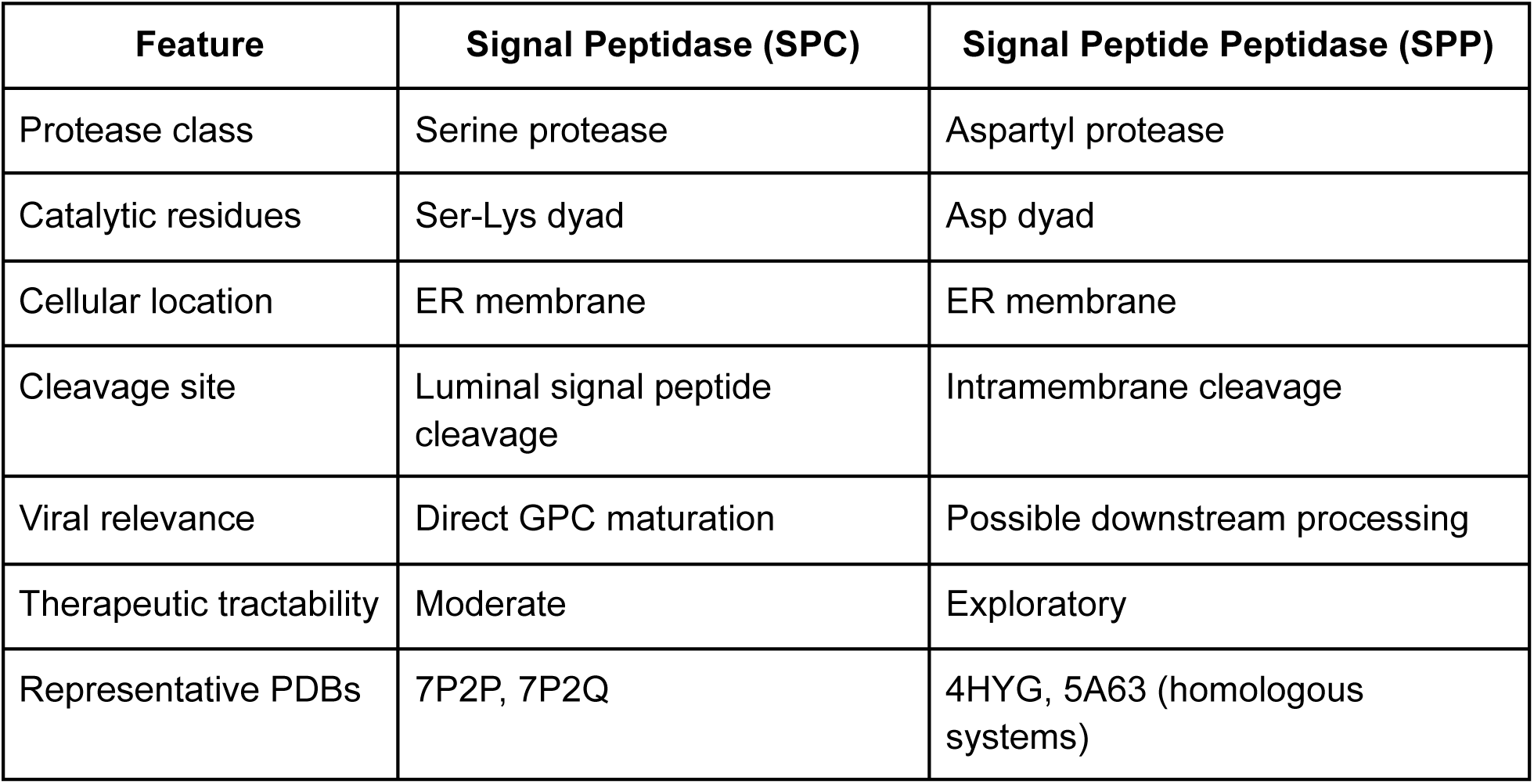
Structural, biochemical, and functional comparison between the signal peptidase complex (SPC) and signal peptide peptidase (SPP) in the context of orthohantavirus glycoprotein processing and host–virus interactions.

| Feature | Signal Peptidase (SPC) | Signal Peptide Peptidase (SPP) |
| --- | --- | --- |
| Protease class | Serine protease | Aspartyl protease |
| Catalytic residues | Ser-Lys dyad | Asp dyad |
| Cellular location | ER membrane | ER membrane |
| Cleavage site | Luminal signal peptide cleavage | Intramembrane cleavage |
| Viral relevance | Direct GPC maturation | Possible downstream processing |
| Therapeutic tractability | Moderate | Exploratory |
| Representative PDBs | 7P2P, 7P2Q | 4HYG, 5A63 (homologous systems) |

## 4. Potential Protease Inhibitor Drugs

To identify potential inhibitors targeting Signal Peptidase I (SP) and Signal Peptide Peptidase (SPP), we queried BRENDA (BRaunschweig ENzyme DAtabase) (Hauenstein et al., 2026), which is currently the most comprehensive online repository of functional, biochemical, and molecular biological data related to enzymes, metabolites, and metabolic pathways. The database was screened for compounds reported as inhibitors of SP and SPP enzymes. Following removal of duplicates, a total of 53 unique compounds were identified for SP, including only two compounds reported for Homo sapiens, whereas 12 unique compounds were identified for SPP, including six compounds associated with Homo sapiens. Consequently, eight compounds were initially considered as potential drug candidates.

To further evaluate their suitability for downstream computational analyses, the identified candidates were cross-referenced with the PubChem database to obtain their corresponding three-dimensional molecular structures. Structural information was successfully retrieved for two SP inhibitors and four SPP inhibitors, resulting in a final set of six compounds selected for further investigation (Table 2).

**Table 2.**
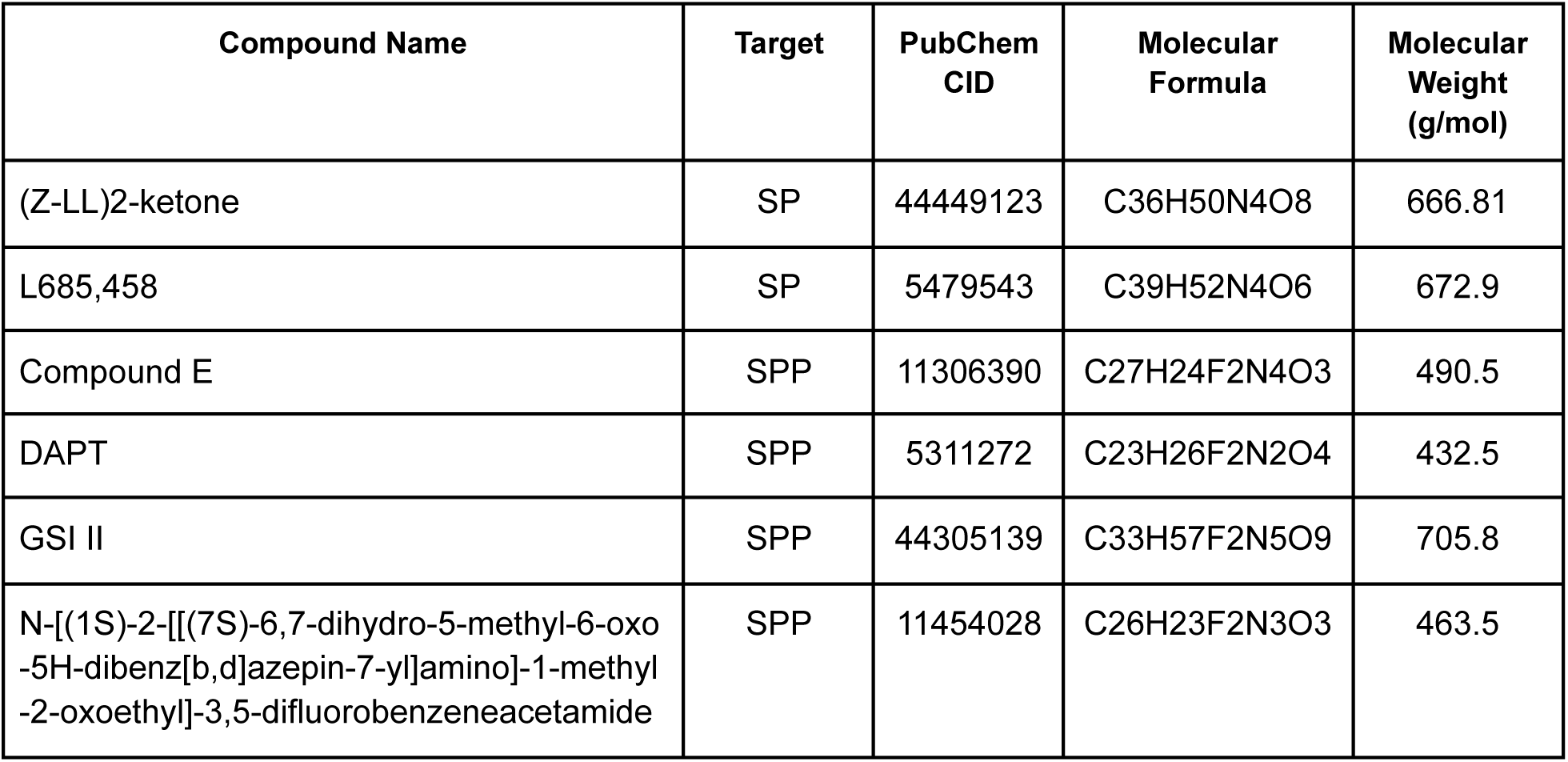
Potential inhibitor compounds identified for Signal Peptidase (SP) and Signal Peptide Peptidase (SPP).

The identified compounds represent structurally diverse protease inhibitors with previously reported activity against intramembrane or signal peptide-processing proteases. Several of these molecules, including DAPT, L685,458, Compound E, and GSI II, are well-known γ-secretase inhibitors, which is consistent with the mechanistic similarity between γ-secretase and Signal Peptide Peptidase, both belonging to the family of intramembrane aspartyl proteases. Their prior characterization in mammalian systems increases their relevance as candidate molecules for SPP inhibition.

Among the identified compounds, Compound E appears particularly promising due to its optimal molecular weight (490.5 g/mol) and superior computational binding characteristics, achieving the highest confidence score (−0.28) in our docking analysis. This difluoro-substituted compound represents an ideal balance between structural complexity and drug-like properties, falling well within Lipinski’s rule of five guidelines while maintaining the sophisticated aromatic scaffold necessary for high-affinity protease binding. Its moderate size and balanced physicochemical profile suggest favorable membrane permeability and bioavailability compared to larger peptide-like inhibitors, while the strategic placement of fluorine atoms may enhance both metabolic stability and binding affinity. The exceptional docking performance of Compound E, combined with its established activity as a γ-secretase inhibitor (Wolfe, 2012), positions it as the most promising lead candidate identified in this study.

Similarly, other difluorophenyl-containing compounds in our dataset, including DAPT (432.5 g/mol), another well-established cell-permeable γ-secretase inhibitor in experimental studies (Dong et al., 2021; Li et al., 2014, Xiao et al., 2014)., exhibit molecular weights below 500 g/mol, collectively representing a class of compounds with favorable drug-like characteristics for further optimization.

In contrast, compounds such as (Z-LL)2-ketone, L685,458, and GSI II possess substantially larger molecular weights exceeding 650 g/mol and exhibit pronounced peptide-like structures. Although such characteristics may confer high inhibitory potency and specificity toward protease active sites, they may also negatively affect oral bioavailability, membrane permeability, and metabolic stability. Peptidomimetic inhibitors are frequently associated with poor pharmacokinetic properties and may require structural optimization or specialized delivery systems for therapeutic application (Gante 1994, Vassiliou et al., 2014).

GSI II, despite being one of the largest compounds identified, contains multiple amide functionalities and fluorinated groups that may contribute to strong target interactions. However, its size and structural complexity could limit its applicability as a conventional small-molecule drug candidate. Similarly, L685,458 is recognized as a potent transition-state analogue inhibitor of γ-secretase (Shearman et al., 2000) but has historically demonstrated limited clinical applicability due to suboptimal pharmacological properties.

Overall, the selected compounds provide a valuable starting point for further computational and experimental evaluation of SP and SPP inhibition. While several candidates demonstrate favorable drug-like characteristics, others primarily serve as high-affinity reference inhibitors that could guide future structure-based optimization efforts. The combination of experimentally validated protease inhibitors and available structural information makes these compounds suitable candidates for subsequent molecular docking, dynamics simulations, and structure-activity relationship analyses.

To further investigate the interaction between selected inhibitors and the targets (SP and SPP), molecular docking analyses were performed using DiffDock (Corso et al., 2023). Compound E was selected as the representative candidate due to its exceptional docking performance, achieving the highest confidence score (−0.28) among all tested compounds, along with favorable drug-like properties including a moderate molecular weight (490.5 g/mol) and balanced physicochemical characteristics (see Supplementary Material for all ligands). Figure 4a presents the molecular structure of Compound E, while Figure 4b illustrates multiple predicted docking conformations generated by DiffDock within the transmembrane binding cavity of Signal Peptide Peptidase. The highest-scoring docking pose (confidence level: −0.28, representing the most favorable binding prediction in our study) is shown in Figure 4c, indicating exceptionally stable localization of the ligand within the putative active-site region. A close-up visualization of the best docking configuration (Figure 4d) reveals that Compound E optimally occupies the hydrophobic pocket formed between transmembrane helices, with its difluoro substitutions and aromatic scaffold providing ideal complementarity for the membrane-embedded catalytic environment. The compact structure allows for efficient packing within the binding cavity while maintaining favorable hydrophobic and electrostatic interactions. These observations, combined with the superior confidence score, strongly support Compound E as the most promising lead compound for structure-based inhibitor optimization targeting SPP.

**Figure 4.**
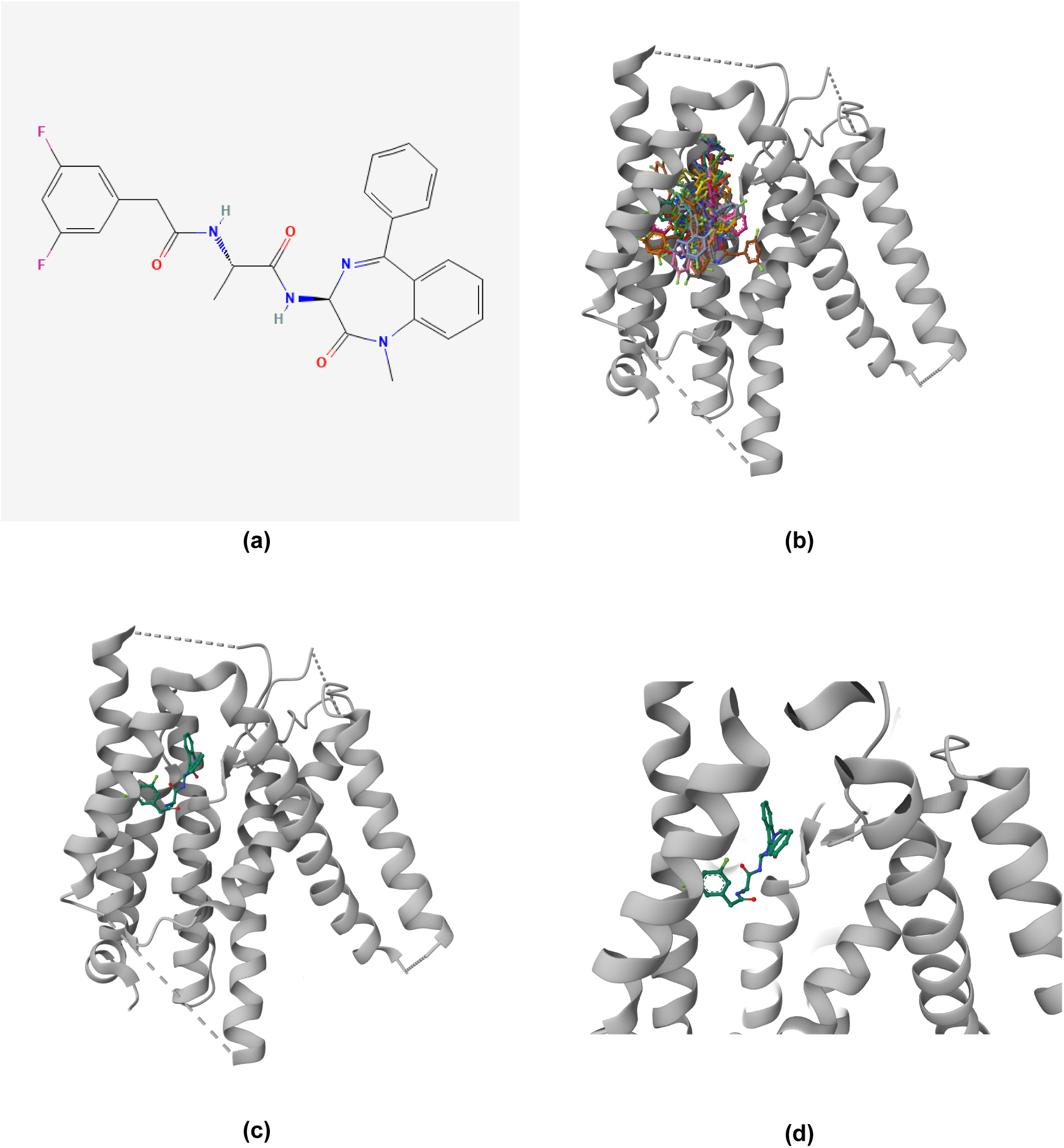
Docking analysis of Compound E with Signal Peptide Peptidase. (a) Molecular structure of Compound E. (b) Multiple predicted docking conformations generated using DiffDock. (c) Highest-scoring docking pose of Compound E within the predicted binding pocket of Signal Peptide Peptidase. (d) Close-up view of the best docking configuration highlighting ligand positioning within the active site.

## 5. Challenges and Future Directions

### 5.1 Current Limitations in Protease-Targeted Antiviral Development

Despite the promising theoretical framework for targeting hantavirus-related proteases, several significant challenges impede the translation of this approach into clinically effective therapeutics. First and foremost, the reliance on host proteases rather than viral enzymes presents a fundamental challenge for selective inhibition. Unlike viral proteases, which represent foreign targets absent from healthy human cells, host proteases such as SPC and SPP perform essential cellular functions. Consequently, broad inhibition of these enzymes could potentially disrupt normal protein processing pathways, leading to cellular toxicity and adverse clinical effects.

The signal peptidase complex, while representing the most mechanistically validated target, processes numerous cellular proteins beyond viral glycoproteins. Inhibition of SPC activity could theoretically interfere with the maturation of secretory proteins, membrane receptors, and other ER-translocated polypeptides essential for normal cellular homeostasis. Similarly, SPP participates in broader ER quality control mechanisms and antigen presentation pathways, suggesting that its inhibition might have unintended immunological consequences.

Our systematic analysis of potential inhibitors revealed additional complexity in the drug development landscape. The identification of only six structurally diverse compounds with appropriate physicochemical properties from the comprehensive BRENDA database highlights the limited chemical space currently available for these targets. The scarcity of compounds is particularly pronounced for SPC inhibitors, with only two suitable candidates identified from human-relevant studies. This finding underscores the need for expanded chemical libraries and novel screening approaches specifically targeting ER-associated proteases.

Furthermore, our molecular docking analysis of DAPT with SPP, while showing promising binding characteristics with a moderate-to-high confidence score (−0.28), represents only preliminary computational validation that requires extensive experimental confirmation. The docking results suggest favorable hydrophobic interactions within the transmembrane catalytic environment, but the translation from computational binding affinity to biological efficacy remains uncertain without biochemical validation studies.

A second major limitation is the current lack of detailed structure-activity relationships (SAR) for hantavirus-related protease inhibitors. While the recent structural characterization of SPC and SPP paralogs has provided valuable architectural insights, the specific molecular determinants governing substrate recognition and cleavage specificity remain incompletely understood. The considerable heterogeneity observed in our inhibitor dataset, with molecular weights ranging from 432.5 to 705.8 g/mol and diverse structural scaffolds including peptidomimetics, small molecules, and difluorophenyl derivatives, underscores the need for systematic SAR studies to optimize drug-like properties while maintaining target selectivity.

The analysis also revealed concerning trends in the physicochemical properties of available inhibitors. Several promising compounds, including (Z-LL)2-ketone, L685,458, and GSI II, exhibit molecular weights exceeding 650 g/mol and pronounced peptide-like characteristics that may negatively impact oral bioavailability, membrane permeability, and metabolic stability. While such compounds may serve as valuable research tools or lead structures, their direct clinical application appears limited without substantial structural optimization.

### 5.2 Therapeutic Window and Selectivity Challenges

The development of effective protease inhibitors must achieve a delicate balance between antiviral efficacy and acceptable toxicity profiles. Given the essential role of host proteases in normal cellular physiology, therapeutic strategies must identify compounds that preferentially disrupt virus-specific protease functions while minimally affecting endogenous protein processing. This selectivity requirement represents a considerably more complex challenge than traditional viral protease inhibition, where complete enzyme suppression is often desirable and achievable without significant host toxicity.

Our computational analysis suggests that achieving such selectivity may be feasible through targeting of virus-specific substrate recognition motifs or allosteric binding sites. The conserved WAASA cleavage motif in hantavirus glycoproteins represents a potential specificity determinant that could be exploited for selective inhibitor design. However, the development of substrate-competitive inhibitors would require detailed understanding of the structural basis for WAASA recognition by SPC, information that remains incomplete despite recent structural advances (Lober et al., 2001).

Furthermore, the temporal dynamics of hantavirus infection may limit the therapeutic window for protease-targeted interventions. Glycoprotein processing occurs early in the viral life cycle, and by the time clinical symptoms become apparent, substantial viral replication and dissemination may have already occurred. This timing constraint suggests that protease inhibitors might be most effective as prophylactic agents in high-risk populations or in combination with other antiviral modalities targeting later stages of infection.

### 5.3 Drug Development and Validation Priorities

Several key research priorities must be addressed to advance protease-targeted antihantavirus therapeutics toward clinical application, informed by our computational analysis and the limitations of currently available inhibitors.

*Enhanced Chemical Library Development*. The limited number of suitable inhibitors identified in our database screening highlights an urgent need for expanded chemical libraries specifically targeting ER-associated proteases. Future screening campaigns should prioritize diversity-oriented synthesis approaches that explore chemical space beyond traditional peptidomimetic scaffolds, particularly focusing on small molecules with favorable ADMET properties.

*Mechanistic Validation Studies*. Rigorous experimental validation of SPC and SPP as bona fide host dependency factors in hantavirus replication remains essential. While biochemical evidence supports the involvement of these proteases in viral glycoprotein maturation, comprehensive loss-of-function studies using the selective inhibitors identified in our analysis (particularly DAPT and related difluorophenyl derivatives) are needed to establish their therapeutic relevance.

*Structure-Guided Inhibitor Optimization*. The recently solved structures of human SPC and SPP paralogs, combined with our docking analysis, provide an opportunity for structure-based lead optimization. Future efforts should focus on improving the drug-like properties of promising candidates such as DAPT while maintaining or enhancing target affinity. This includes addressing the molecular weight and solubility limitations observed in our current inhibitor set.

*Selectivity Profiling and Safety Assessment*. Comprehensive evaluation of inhibitor selectivity across the broader cellular protease landscape will be crucial for identifying compounds with acceptable safety profiles. Our analysis identified several γ-secretase inhibitors (DAPT, L685,458, GSI II) that may exhibit cross-reactivity with related intramembrane proteases, necessitating careful toxicological evaluation.

*Combination Therapy Development*. Given the limitations of single-target approaches highlighted by our analysis, future therapeutic strategies should explore combination regimens incorporating protease inhibitors alongside agents targeting other aspects of hantavirus biology, including viral entry, RNA replication, or host immune responses.

## 6. Conclusion

Orthohantaviruses represent a persistent and evolving global health threat, with recent outbreak events underscoring the urgent need for effective therapeutic interventions. The absence of approved antiviral treatments, combined with the high mortality rates associated with HFRS and HCPS (Khaiboullina et al., 2005) , creates a compelling rationale for innovative drug development approaches. This study has systematically examined the potential of targeting hantavirus-related proteases, particularly host ER proteases involved in viral glycoprotein maturation, as a novel antiviral strategy through comprehensive database mining, structural analysis, and computational modeling.

Our investigation confirms that the dependence of hantaviruses on host proteolytic machinery for essential steps in viral maturation represents both an opportunity and a challenge for therapeutic intervention. The signal peptidase complex (SPC) emerges as the most mechanistically validated target, given its direct role in processing the viral glycoprotein precursor at the conserved WAASA motif. The recent structural characterization of human SPC paralogs provides a solid foundation for structure-based inhibitor design, while the essential nature of this processing step suggests that selective inhibition could effectively disrupt viral replication.

Signal peptide peptidase (SPP) represents a more exploratory but potentially valuable target, given its role in downstream processing of membrane-embedded peptide fragments and its involvement in broader ER quality control mechanisms. Our molecular docking analysis of Compound E with SPP yielded exceptional results (confidence level: −0.28, the highest among all tested compounds), strongly suggesting that this γ-secretase inhibitor represents the most promising lead compound for further optimization. The structural insights gained from this computational analysis provide a foundation for rational drug design efforts targeting this intramembrane protease.

However, our systematic analysis also reveals significant challenges that must be addressed for successful clinical translation. The identification of only six suitable inhibitors from comprehensive database screening highlights the limited chemical space currently available for these targets, with particularly concerning gaps in SPC inhibitor development. The predominance of large, peptide-like molecules among available inhibitors raises substantial concerns about drug-like properties, bioavailability, and clinical feasibility.

The pursuit of host protease-targeted therapies faces fundamental challenges that distinguish this approach from traditional viral enzyme inhibition. The dual role of host proteases in both viral and cellular processes demands the development of highly selective inhibitors capable of discriminating between pathological and physiological substrate processing. Our analysis of available inhibitors reveals significant molecular weight and structural diversity challenges that must be overcome to achieve optimal therapeutic profiles.

Despite these obstacles, several factors support the continued investigation of protease-targeted antihantavirus therapeutics. The conserved nature of host proteolytic machinery across different hantavirus species suggests that successful inhibitors could provide broad-spectrum activity against both Old and New World viruses. Furthermore, the essential role of glycoprotein processing in viral infectivity, confirmed by our mechanistic analysis, implies that even partial disruption of protease activity might yield significant antiviral benefits.

Our computational modeling results, while preliminary, provide proof-of-concept evidence that selective targeting of these proteases is feasible. The moderate-to-high confidence binding prediction for Compound E suggests that optimization of existing γ-secretase inhibitors could yield compounds with enhanced specificity for viral substrates while minimizing cellular toxicity.

The integration of structural biology advances, computational design approaches, and systematic inhibitor profiling demonstrated in this study positions the field for meaningful progress in protease-targeted drug development. Our analysis identifies specific research priorities, including expanded chemical library development, structure-guided lead optimization, and comprehensive selectivity profiling, that should guide future investigations.

In the broader context of emerging infectious disease preparedness, the development of host-directed antiviral therapies represents a valuable complement to traditional pathogen-specific approaches. While the challenges associated with targeting essential cellular processes are substantial, as highlighted by our analysis, the potential for broad-spectrum activity and reduced likelihood of viral resistance development make host protease inhibition an attractive long-term strategy.

The recent hantavirus outbreak serves as a stark reminder of the ongoing threat posed by these pathogens and the critical need for effective therapeutic options. Our systematic analysis provides a roadmap for addressing the key challenges in protease-targeted drug development while identifying specific molecular targets and lead compounds for further investigation.

Ultimately, the successful translation of protease-targeted antihantavirus therapeutics from concept to clinic will require sustained research investment, innovative technological approaches, and careful attention to the unique challenges posed by host-directed therapy. Our study provides a foundation for these efforts by systematically characterizing the available chemical space, identifying promising lead compounds, and highlighting key research priorities for future investigation.

The path forward demands continued collaboration between virologists, structural biologists, medicinal chemists, and clinicians to address the multifaceted challenges of targeting host proteases for antiviral therapy. The computational and structural insights generated in this study provide a valuable starting point for such collaborative efforts, offering specific molecular targets, lead compounds, and optimization strategies that can guide future drug development initiatives.

## Materials and Methods

### Database Mining and Inhibitor Identification

Potential protease inhibitors targeting Signal Peptidase (EC 3.4.21.89) and Signal Peptide Peptidase (EC 3.4.23.B24) were systematically identified through comprehensive screening of the BRENDA (BRaunschweig ENzyme DAtabase, https://www.brenda-enzymes.org), which serves as the most comprehensive online repository of functional, biochemical, and molecular biological data related to enzymes, metabolites, and metabolic pathways. The database query was implemented in Python using automated web scraping protocols to retrieve all reported inhibitors for the target enzymes.

Following initial data collection, duplicate entries were removed through automated comparison of compound names and chemical identifiers. Species-specific filtering was applied to prioritize compounds with reported activity in Homo sapiens systems. The resulting candidate compounds were cross-referenced with the PubChem database to obtain three-dimensional molecular structures and physicochemical properties.

### Structural Data Acquisition and Processing

Three-dimensional protein structures for human Signal Peptidase Complex paralogs (PDB IDs: 7P2P, 7P2Q) and Signal Peptide Peptidase-Like 2A (PDB IDs: 9K92, 9K93) were downloaded from the Protein Data Bank. Structural processing and visualization were performed using Python-based molecular modeling libraries, including BioPython and PyMOL API integration.

### Chemical Structure Processing

Small molecule structures were obtained in SMILES format from PubChem and converted to three-dimensional SDF format using the NCI CACTUS Chemical Identifier Resolver (https://cactus.nci.nih.gov/translate). All chemical structure manipulations, molecular property calculations, and data processing were implemented in Python using RDKit and related cheminformatics libraries.

### Molecular Docking Analysis

Molecular docking studies were performed using DiffDock (https://build.nvidia.com/mit/diffdock), a state-of-the-art deep learning-based molecular docking platform. Docking parameters were set to generate 20 poses per ligand with 18 diffusion steps and 20 time divisions to ensure comprehensive sampling of binding configurations.

Docking confidence scores were interpreted according to the following criteria (Corso et al., 2023):

● Score > 0: High confidence (strong prediction, likely accurate binding pose)
● Score −1.5 to 0: Moderate confidence (reasonable prediction, may need validation)
● Score < −1.5: Low confidence (uncertain prediction, requires careful validation)

### Computational Infrastructure and Data Analysis

All computational analyses, data processing, visualization, and statistical calculations were implemented in Python using standard scientific computing libraries including NumPy, Pandas, Matplotlib, and Seaborn. Molecular structure analysis and manipulation were performed using RDKit and Open Babel integration. Custom analysis scripts were developed for automated database querying, structure-activity relationship analysis, and results visualization.

The complete computational workflow, from database mining through molecular docking analysis, was designed to ensure reproducibility and scalability for future expansion to additional protease targets and larger chemical libraries.

### External Tool Integration

Two external web-based tools were incorporated into the analysis pipeline: the NCI CACTUS Chemical Identifier Resolver for SMILES to 3D structure conversion, and the NVIDIA DiffDock platform for molecular docking calculations. All other computational tasks, including data preprocessing, analysis, and visualization, were implemented using Python-based solutions to maintain consistency and reproducibility across the entire workflow.

## Authors’ Contributions

Conceptualization: EWT; Methodology: EWT, JMT; Visualization: JMT; Writing (Original Draft): JMT; Writing (Review): EWT; Software: JMT; Supervision: EWT

---

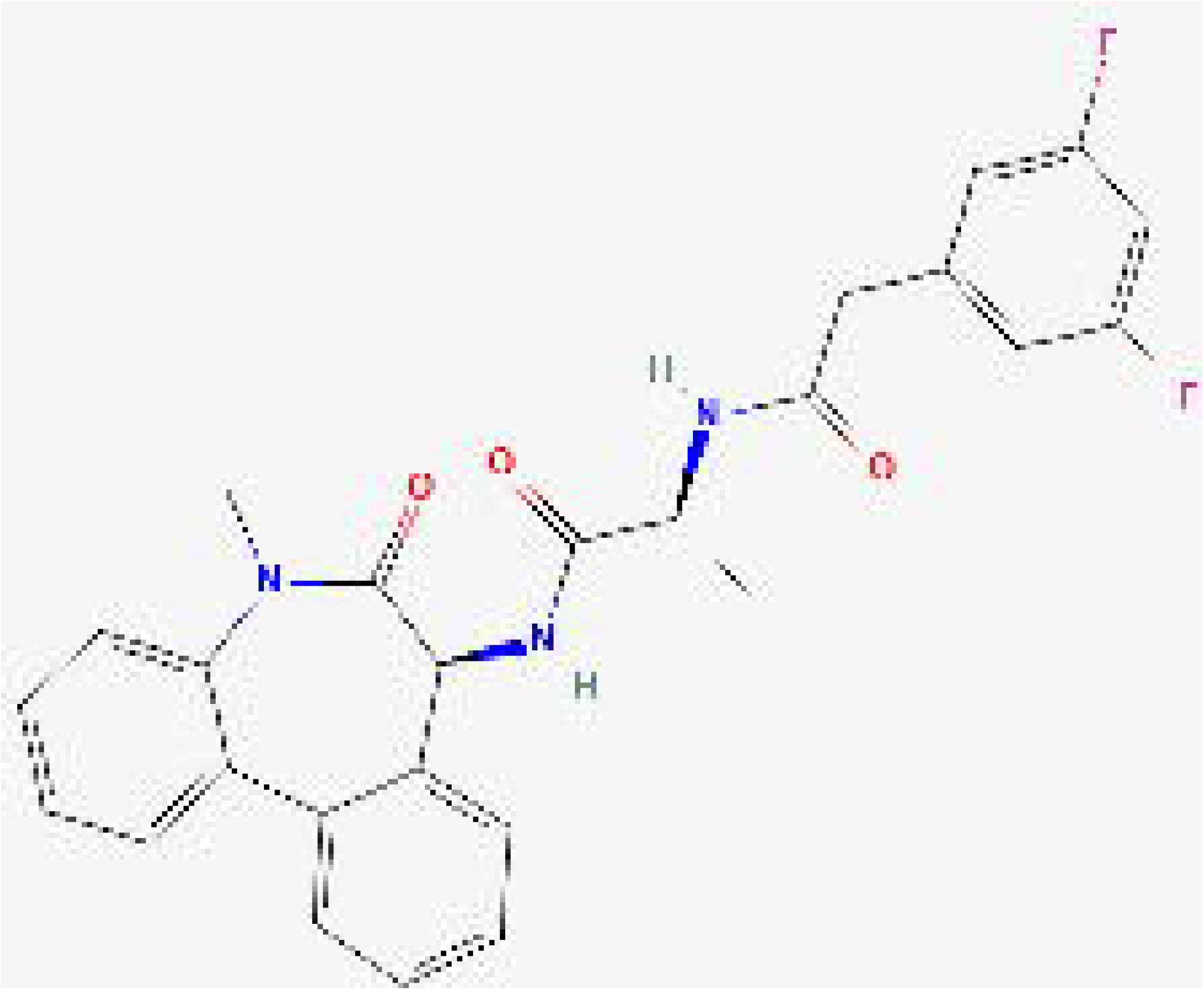

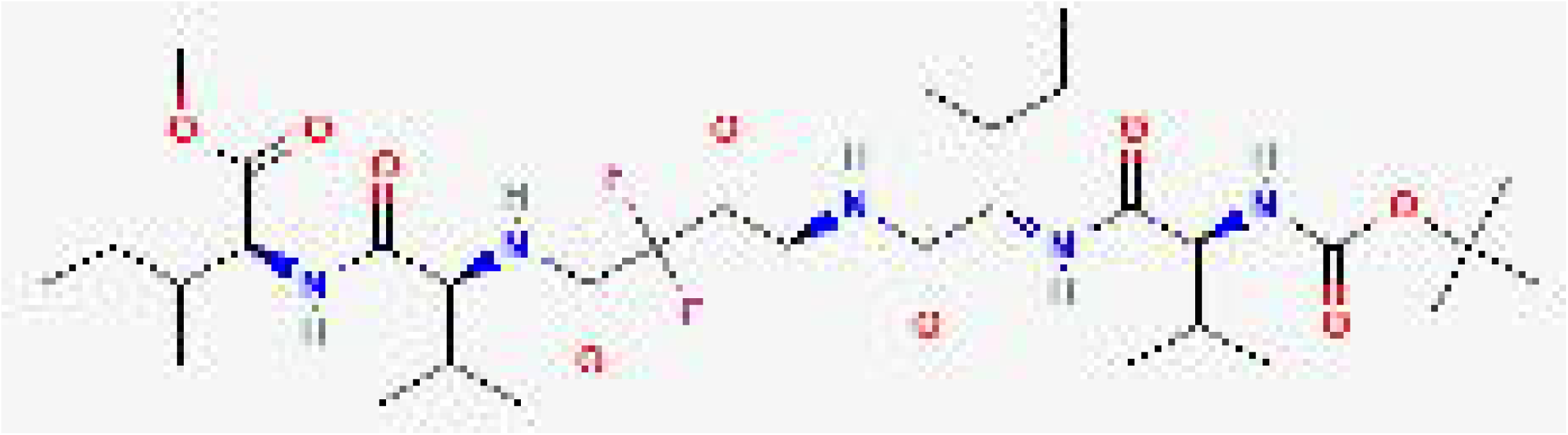

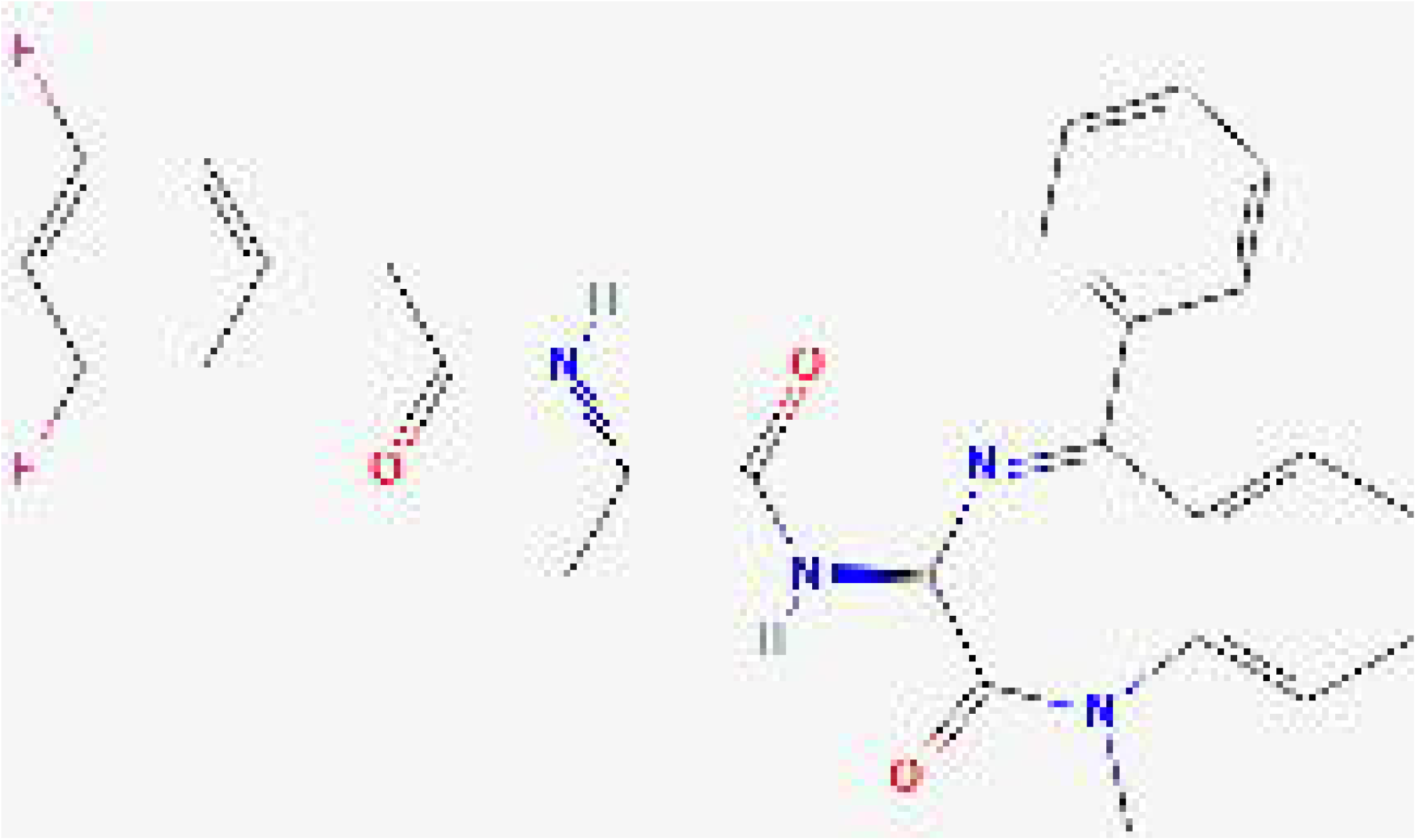

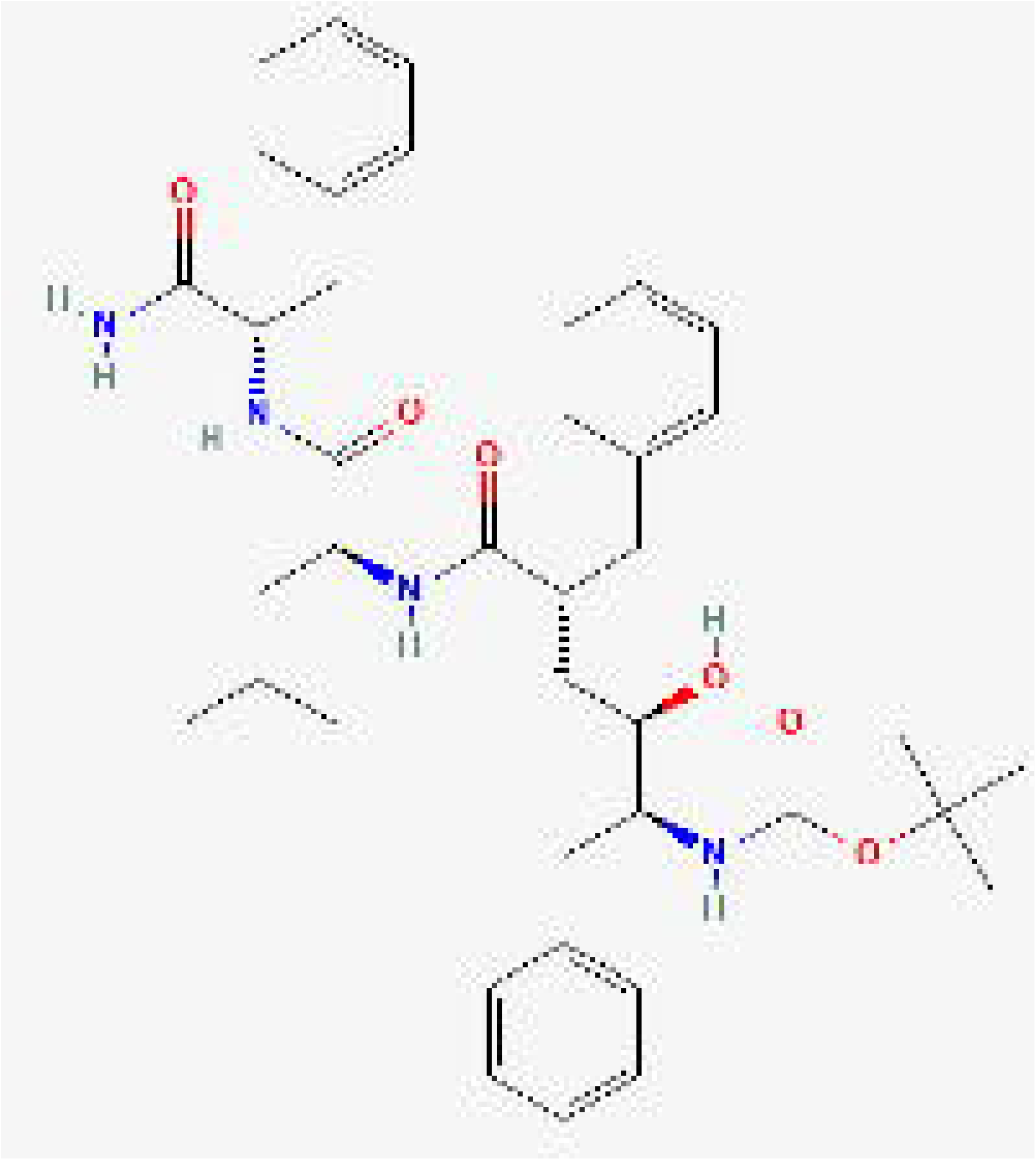

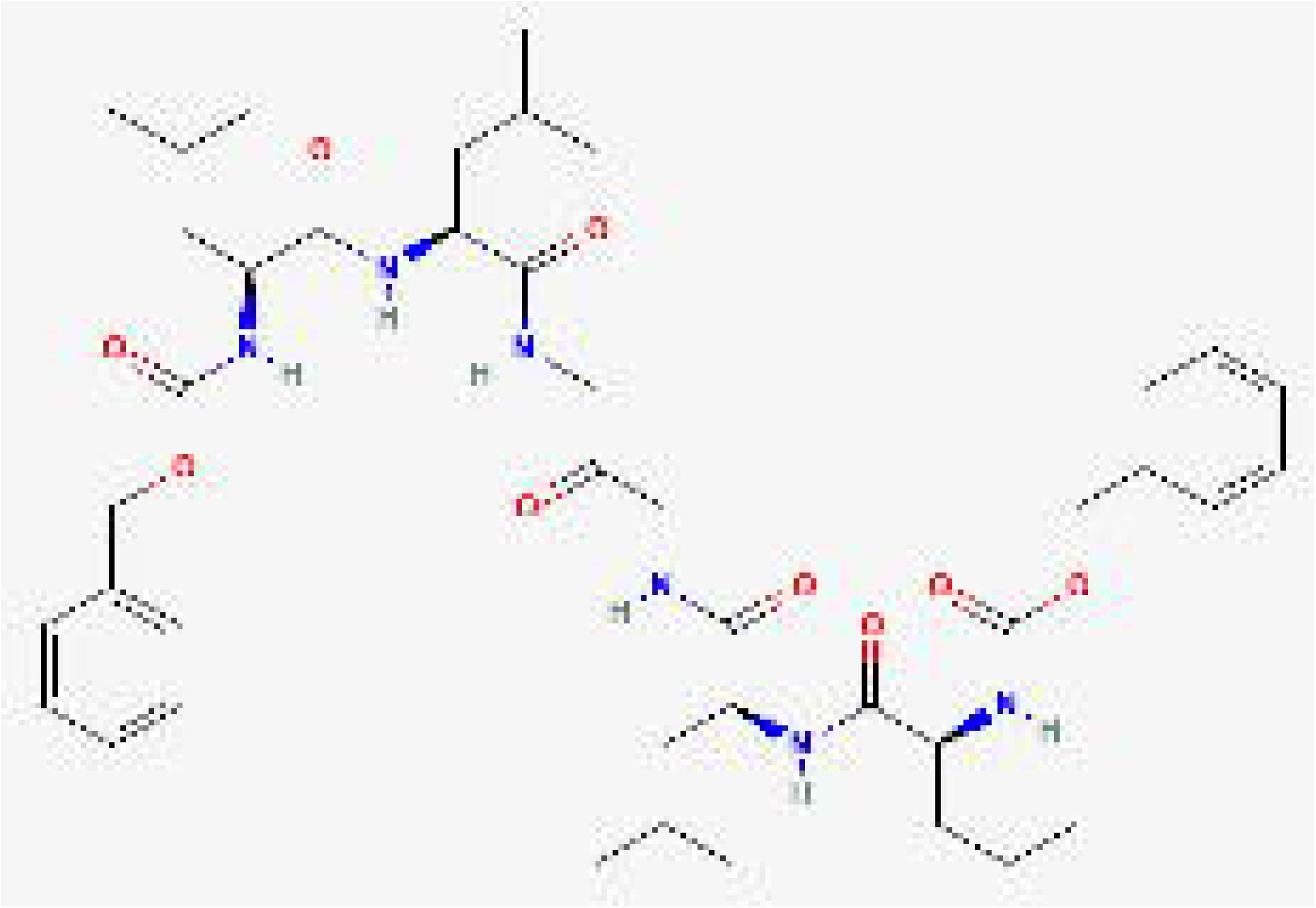

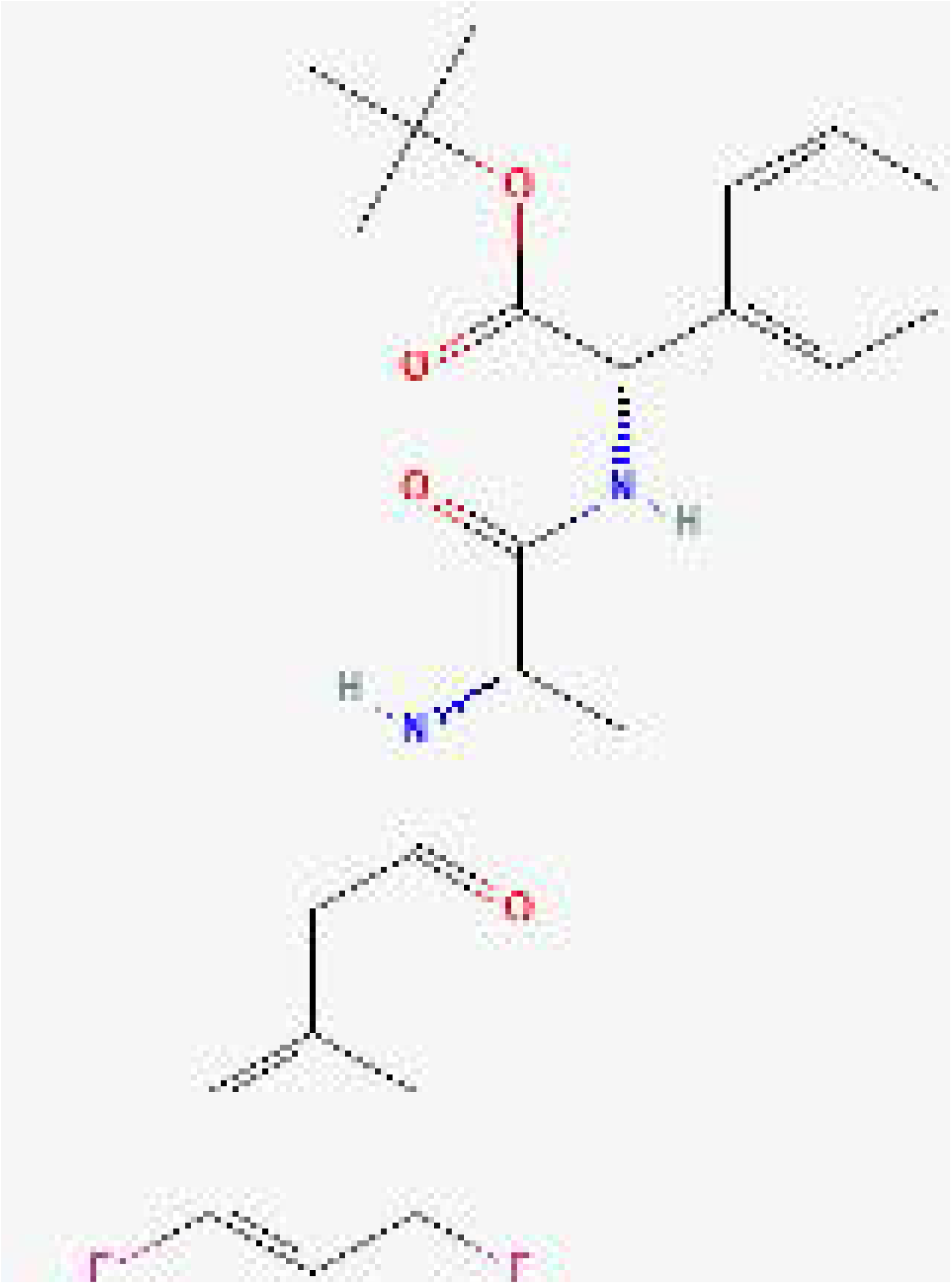

## References

(Acuna, et al., 2014) Acuña, Rodrigo, et al. “Hantavirus Gn and Gc glycoproteins self-assemble into virus-like particles.” Journal of virology 88.4 (2014): 2344-2348.

(Afzal, et al., 2023) Afzal, Samia, et al. “Hantavirus: an overview and advancements in therapeutic approaches for infection.” Frontiers in Microbiology 14 (2023): 1233433.

(Agbowuro, et al., 2018) Agbowuro, Ayodeji A., et al. “Proteases and protease inhibitors in infectious diseases.” Medicinal research reviews 38.4 (2018): 1295-1331.

(Auclair, et al., 2012) Auclair, S. M., Bhanu, M. K., & Kendall, D. A. (2012). Signal peptidase I: cleaving the way to mature proteins. Protein Science, 21(1), 13–25.

(Brix, 2018) Brix, Klaudia. “Host cell proteases: Cathepsins.” Activation of viruses by host proteases. Cham: Springer International Publishing, 2018. 249–276.

(Cifuentes-Munoz, et al., 2014) Cifuentes-Muñoz, Nicolás, Natalia Salazar-Quiroz, and Nicole D. Tischler. “Hantavirus Gn and Gc envelope glycoproteins: key structural units for virus cell entry and virus assembly.” Viruses 6.4 (2014): 1801-1822.

(Corso et al., 2023) Corso, Gabriele, et al. “DiffDock: Diffusion Steps, Twists, and Turns for Molecular Docking.” International Conference on Learning Representations (ICLR 2023). 2023.

(Dong et al., 2021) Dong, Zimei, et al. “Gamma-Secretase Inhibitor (DAPT), a potential therapeutic target drug, caused neurotoxicity in planarian regeneration by inhibiting Notch signaling pathway.” Science of the Total Environment 781 (2021): 146735.

(Dunn, et al., 2002) Dunn, Ben M., et al. “Retroviral proteases.” Genome biology 3.4 (2002): reviews3006-1

(Gante 1994) Gante, Joachim. “Peptidomimetics—tailored enzyme inhibitors.” Angewandte Chemie International Edition in English 33.17 (1994): 1699-1720.

(Hart & Bennett, 1999) Hart, C. A., and M. Bennett. “Hantavirus infections: epidemiology and pathogenesis.” Microbes and infection 1.14 (1999): 1229-1237.

(Hauenstein et al., 2026) Hauenstein, Julia, et al. “BRENDA in 2026: a Global Core Biodata Resource for functional enzyme and metabolic data within the DSMZ Digital Diversity.” Nucleic Acids Research 54.D1 (2026): D527–D534.

(Huag, et al., 2025) Huang, Gaoxingyu, et al. “Structural insights into human signal peptide peptidase.” Proceedings of the National Academy of Sciences 122.51 (2025): e2528340122.

(Khaiboullina, et al., 2005) Khaiboullina, Svetlana F., S. P. Morzunov, and Stephen C. St Jeor. “Hantaviruses: Molecular biology, evolution and pathogenesis.” Current molecular medicine 5.8 (2005): 773-790.

(Lemberg, et al., 2002) Lemberg, Marius K., and Bruno Martoglio. “Requirements for signal peptide peptidase-catalyzed intramembrane proteolysis.” Molecular cell 10.4 (2002): 735-744.

(Li, et al., 2014) Li, Jun-yang, Ru-jun Li, and Han-dong Wang. “γ-secretase inhibitor DAPT sensitizes t-AUCB-induced apoptosis of human glioblastoma cells in vitro via blocking the p38 MAPK/MAPKAPK2/Hsp27 pathway.” Acta Pharmacologica Sinica 35.6 (2014): 825–831.

(Liaci, et al., 2021) Liaci, A. Manuel, et al. “Structure of the human signal peptidase complex reveals the determinants for signal peptide cleavage.” Molecular cell 81.19 (2021): 3934-3948.

(Lober, et al., 2001) Löber, Christian, et al. “The Hantaan virus glycoprotein precursor is cleaved at the conserved pentapeptide WAASA.” Virology 289.2 (2001): 224-229.

(Meier, et al., 2021) Meier, Kristina, et al. “Hantavirus replication cycle—an updated structural virology perspective.” Viruses 13.8 (2021): 1561.

(Muyangwa, et al., 2015) Muyangwa, Musalwa, et al. “Hantaviral proteins: structure, functions, and role in hantavirus infection.” Frontiers in microbiology 6 (2015): 1326.

(Paetzel, et al., 2002) Paetzel, Mark, et al. “Signal peptidases.” Chemical reviews 102.12 (2002): 4549-4580.

(Schwake, et al., 2022) Schwake, Christopher, Michael Hyon, and Athar H. Chishti. “Signal peptide peptidase: a potential therapeutic target for parasitic and viral infections.” Expert opinion on therapeutic targets 26.3 (2022): 261-273.

(Senkowska & Weglarz-Tomczak, 2026) Senkowska, Zuzanna, and Ewelina Weglarz-Tomczak. “Decoding the Secretase Puzzle in Amyloid-β Generation: A State-of-the-Art Overview of the Protease-Mediated APP Processing Cascade in Alzheimer’s Disease.” Ageing Research Reviews (2026): 10311

(Shearman, et al., 2000) Shearman, Mark S., et al. “L-685,458, an aspartyl protease transition state mimic, is a potent inhibitor of amyloid β-protein precursor γ-secretase activity.” Biochemistry 39.30 (2000): 8698–8704.

(Shi, et al., 2016) Shi, Xiaohong, et al. “Bunyamwera orthobunyavirus glycoprotein precursor is processed by cellular signal peptidase and signal peptide peptidase.” Proceedings of the National Academy of Sciences 113.31 (2016): 8825-8830.

(Teramoto et al., 2023) Teramoto, Tadahisa, Kyung H. Choi, and Radhakrishnan Padmanabhan. “Flavivirus proteases: The viral Achilles heel to prevent future pandemics.” Antiviral research 210 (2023): 105516.

(Turk, 2006) Turk, Boris. “Targeting proteases: successes, failures and future prospects.” Nature reviews Drug discovery 5.9 (2006): 785-799.

(Vassiliou, et al., 2014) Vassiliou, Stamatia, et al. “Structure-guided, single-point modifications in the phosphinic dipeptide structure yield highly potent and selective inhibitors of neutral aminopeptidases.” Journal of medicinal chemistry 57.19 (2014): 8140-8151.

(Voss, et al. 2013) Voss, Matthias, Bernd Schröder, and Regina Fluhrer. “Mechanism, specificity, and physiology of signal peptide peptidase (SPP) and SPP-like proteases.” Biochimica et Biophysica Acta (BBA)-Biomembranes 1828.12 (2013): 2828-2839.

(Weihofen, et al., 2002) Weihofen, Andreas, et al. “Identification of signal peptide peptidase, a presenilin-type aspartic protease.” Science 296.5576 (2002): 2215-2218.

(Weglarz-Tomczak, et al., 2018) Weglarz-Tomczak, Ewelina, et al. “Neutral metalloaminopeptidases APN and MetAP2 as newly discovered anticancer molecular targets of actinomycin D and its simple analogs.” Oncotarget 9.50 (2018): 29365.

(Weglarz-Tomczak, et al., 2021) Weglarz-Tomczak, Ewelina, et al. “Identification of ebselen and its analogues as potent covalent inhibitors of papain-like protease from SARS-CoV-2.” Scientific reports 11.1 (2021): 3640.

(Wolfe, 2012) Wolfe, Michael S. “γ-Secretase inhibitors and modulators for Alzheimer’s disease.” Journal of neurochemistry 120 (2012): 89–98.

(Xiao, et al., 2014) Xiao, Yong Guang, et al. “γ-Secretase inhibitor DAPT attenuates intimal hyperplasia of vein grafts by inhibition of Notch1 signaling.” Laboratory investigation 94.6 (2014): 654–662.

(Zanotti, et al., 2022) Zanotti, Andrea, et al. “The human signal peptidase complex acts as a quality control enzyme for membrane proteins.” Science 378.6623 (2022): 996-1000.

