## Supplementary Material for "Orthohantavirus-related Proteases as Therapeutic Targets: Opportunities for Antiviral Drug Development"

#### 1. Identified potential inhibitors of Signal Peptidase (SP) and Signal Peptide Peptidase (SPP)

The analysis carried out in the paper resulted in a final set of six compounds selected for further investigation (Table S1).

**Table S1. Potential inhibitor compounds identified for Signal Peptidase (SP) and Signal Peptide Peptidase (SPP).**

| Compound Name | Target | PubChem CID | Molecular Formula | Molecular Weight (g/mol) |
|---|---|---|---|---|
| (Z-LL)2-ketone | SP | 44449123 | C36H50N4O8 | 666.81 |
| L685,458 | SP | 5479543 | C39H52N4O6 | 672.9 |
| Compound E | SPP | 11306390 | C27H24F2N4O3 | 490.5 |
| DAPT | SPP | 5311272 | C23H26F2N2O4 | 432.5 |
| GSI II | SPP | 44305139 | C33H57F2N5O9 | 705.8 |
| N-[(1S)-2-[[(7S)-6,7-dihydro-5-methyl-6-oxo-5H-dibenz[b,d]azepin-7-yl]amino]-1-methyl-2-oxoethyl]-3,5-difluorobenzeneacetamide | SPP | 11454028 | C26H23F2N3O3 | 463.5 |

### 2. Molecular docking

To further investigate the interaction between selected inhibitors and the targets (SP and SPP), molecular docking analyses were performed using DiffDock (Corso et al., 2023). In the following, we present results of molecular docking for all ligands, and provide the confidence levels for the best candidates for each inhibitor.

#### Signal Peptidase

**(Z-LL)2-ketone.** The molecular docking analysis of (Z-LL)2-ketone with Signal Peptidase I reveals a moderate binding interaction with a confidence level of -2.05. The docking results show that this dipeptide ketone inhibitor successfully occupies the predicted binding pocket of Signal Peptidase I. The molecular structure displays the characteristic Z-protected leucyl-leucine backbone with ketone functionality, which is designed to mimic the transition state of peptide hydrolysis. The binding pose indicates that the compound fits well within the active site cavity, with the ketone group potentially positioned to interact with the catalytic residues. The multiple conformations generated by DiffDock suggest some flexibility in binding orientation, but the highest-scoring pose shows stable positioning within the enzyme's active site. See Figure S1.

**L685,458.** L685,458 demonstrates binding to Signal Peptidase I with a confidence level of -2.30, representing the lowest confidence among all tested compounds. This peptidomimetic inhibitor shows a more extended conformation compared to (Z-LL)2-ketone, with multiple aromatic rings and peptide-like linkages. The docking analysis reveals that despite its larger size and structural complexity, L685,458 can accommodate within the SP I binding pocket. The molecular visualization shows the compound adopting a conformation that allows multiple interaction points with the enzyme. However, the relatively low confidence score suggests that this binding mode may be less favorable or stable compared to other candidates. See Figure S2.

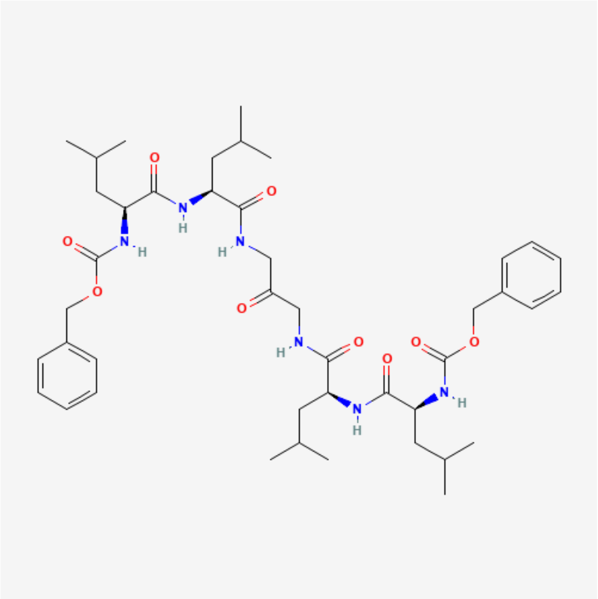

(a)

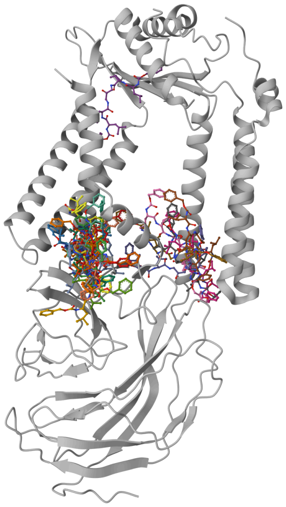

(b)

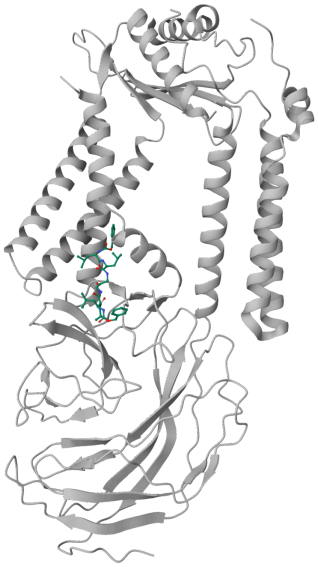

(c)

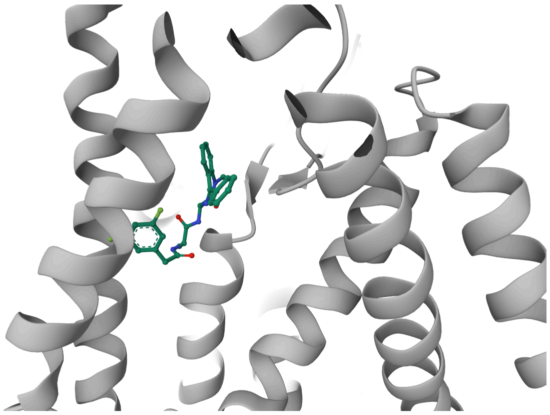

(d)

**Figure S1. Docking analysis of (Z-LL)2-ketone with Signal Peptidase I.**

(a) Molecular structure of (Z-LL)2-ketone. (b) Multiple predicted docking conformations generated using DiffDock. (c) Highest-scoring docking pose of (Z-LL)2-ketone within the predicted binding pocket of Signal Peptidase I. (d) Close-up view of the best docking configuration highlighting ligand positioning within the active site.

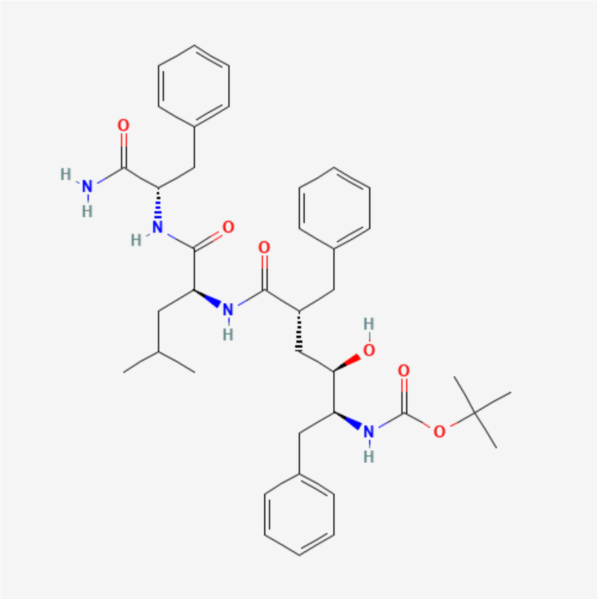

**(a)**

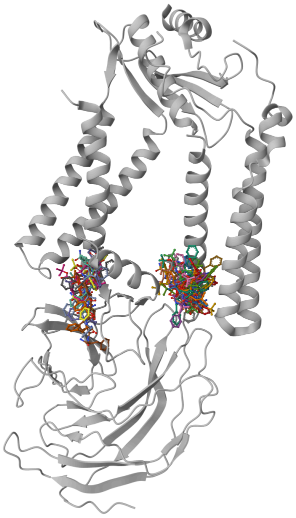

**(b)**

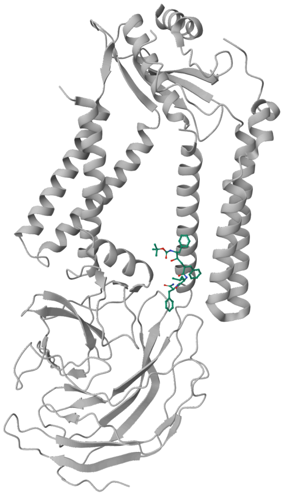

**(c)**

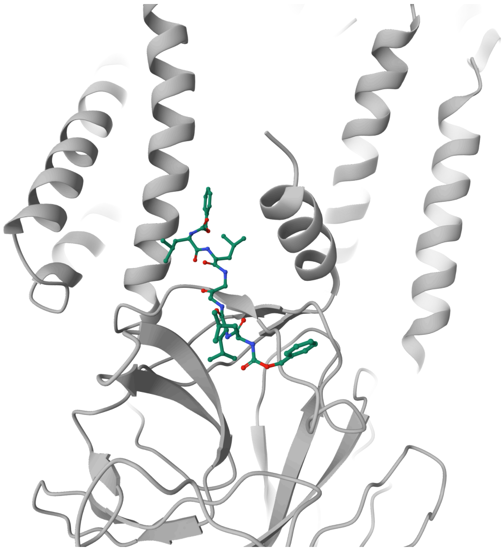

**(d)**

**Figure S2. Docking analysis of L685,458 with Signal Peptidase I.**
(a) Molecular structure of L685,458. (b) Multiple predicted docking conformations generated using DiffDock. (c) Highest-scoring docking pose of L685,458 within the predicted binding pocket of Signal Peptidase I. (d) Close-up view of the best docking configuration highlighting ligand positioning within the active site.

#### Signal Peptide Peptidase

**Compound E.** Compound E exhibits promising binding to Signal Peptide Peptidase with a confidence level of -0.28, which is the highest (most favorable) confidence score among all tested compounds. This smaller molecule with difluoro substitution shows excellent complementarity with the SPP binding pocket. The docking results indicate that Compound E achieves a well-defined binding pose within the active site, suggesting strong potential for inhibitory activity. The compact structure and optimal positioning within the binding cavity, combined with the highest confidence level, make this compound particularly attractive for further development as an SPP inhibitor. It is treated as a classic high-potency research γ-secretase inhibitor that helped define the field, but was superseded clinically by safer and more optimized molecules. See Figure S3.

**DAPT.** DAPT is the smallest molecule from all six inhibitors, and shows binding to Signal Peptide Peptidase with a confidence level of -1.11, representing moderate binding affinity. This well-known γ-secretase inhibitor contains difluorobenzyl and diphenylacetyl groups that appear to interact favorably with the SPP binding site. The docking visualization shows DAPT fitting well within the active site cavity, with the aromatic rings potentially forming beneficial hydrophobic interactions. The moderate confidence level suggests that while DAPT can bind to SPP, the interaction may not be as optimal as with Compound E, but still represents a viable inhibitory interaction. See Figure S4.

**GSI II.** GSI II demonstrates binding to Signal Peptide Peptidase with a confidence level of -2.03, indicating relatively weak binding among the SPP inhibitors tested. This larger, more complex molecule with difluoro substitution and extended peptidic structure shows the ability to fit within the SPP binding pocket, though with lower confidence than the other SPP-targeted compounds. The docking results suggest that while GSI II can achieve binding, the interaction may be less stable or optimal compared to Compound E and DAPT. The complex structure may lead to some steric hindrance or suboptimal positioning within the active site. See Figure S5.

**N-[(1S)-2-[[(7S)-6,7-dihydro-5-methyl-6-oxo-5H-dibenz[b,d]azepin-7-yl]amino]-1-methyl-2-oxoethyl]-3,5-difluorobenzeneacetamide.** This structurally complex compound shows binding to Signal Peptide Peptidase with a confidence level of -0.99, representing good binding potential. The molecule contains a dibenzazepine core with difluorobenzene substitution, creating a substantial binding interface with the SPP active site. The docking analysis reveals that despite its size and complexity, the compound can achieve favorable positioning within the binding pocket. The confidence level suggests moderate to good binding affinity, making it a reasonable candidate for SPP inhibition, though not as promising as Compound E based purely on the docking confidence scores. See Figure S6.

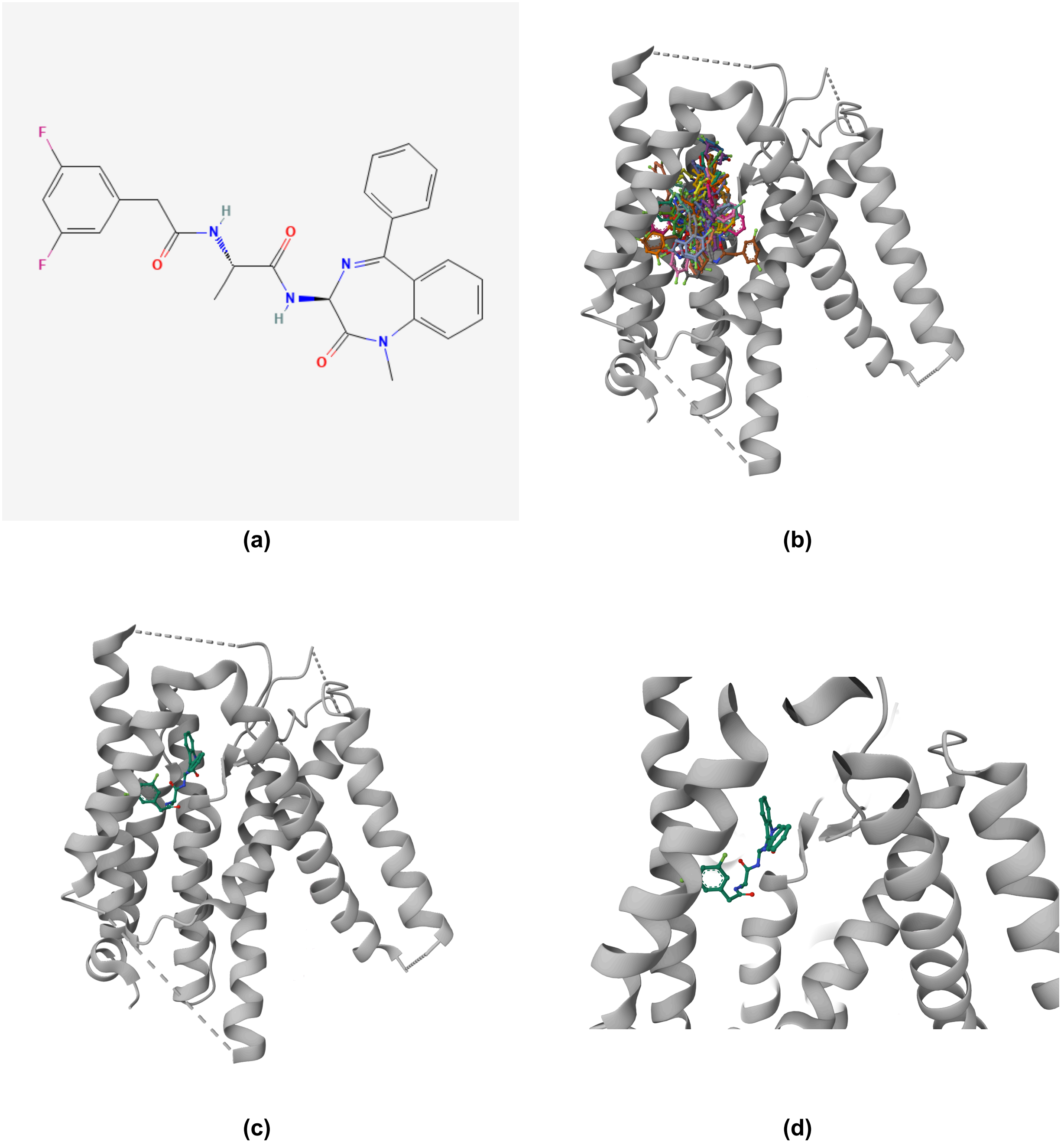


**Figure S3. Docking analysis of Compound E with Signal Peptide Peptidase.**
(a) Molecular structure of Compound E. (b) Multiple predicted docking conformations generated using DiffDock. (c) Highest-scoring docking pose of Compound E within the predicted binding pocket of Signal Peptide Peptidase. (d) Close-up view of the best docking configuration highlighting ligand positioning within the active site.

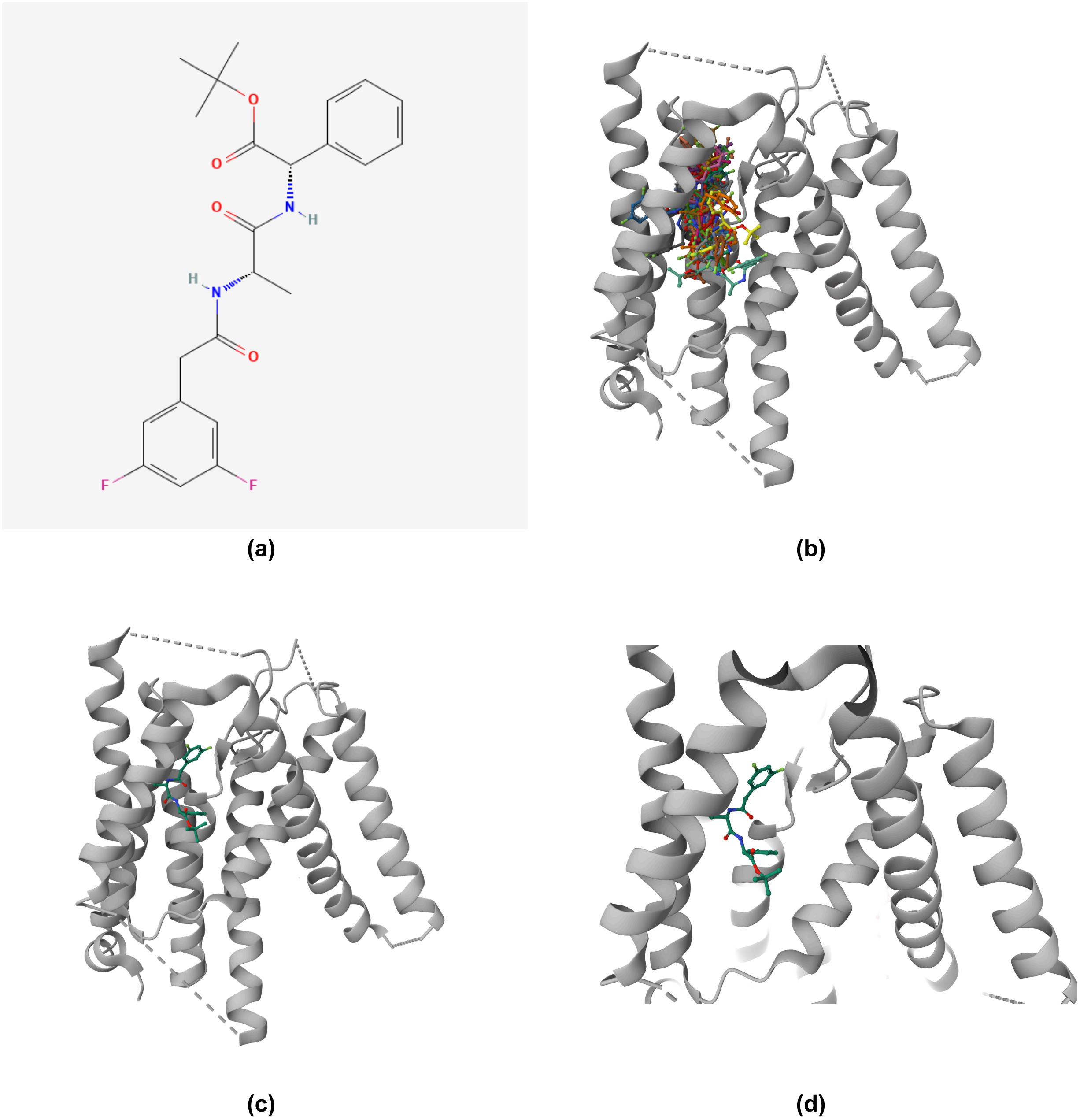


**Figure S4. Docking analysis of DAPT with Signal Peptide Peptidase.**
(a) Molecular structure of DAPT. (b) Multiple predicted docking conformations generated using DiffDock. (c) Highest-scoring docking pose of DAPT within the predicted binding pocket of Signal Peptide Peptidase. (d) Close-up view of the best docking configuration highlighting ligand positioning within the active site.

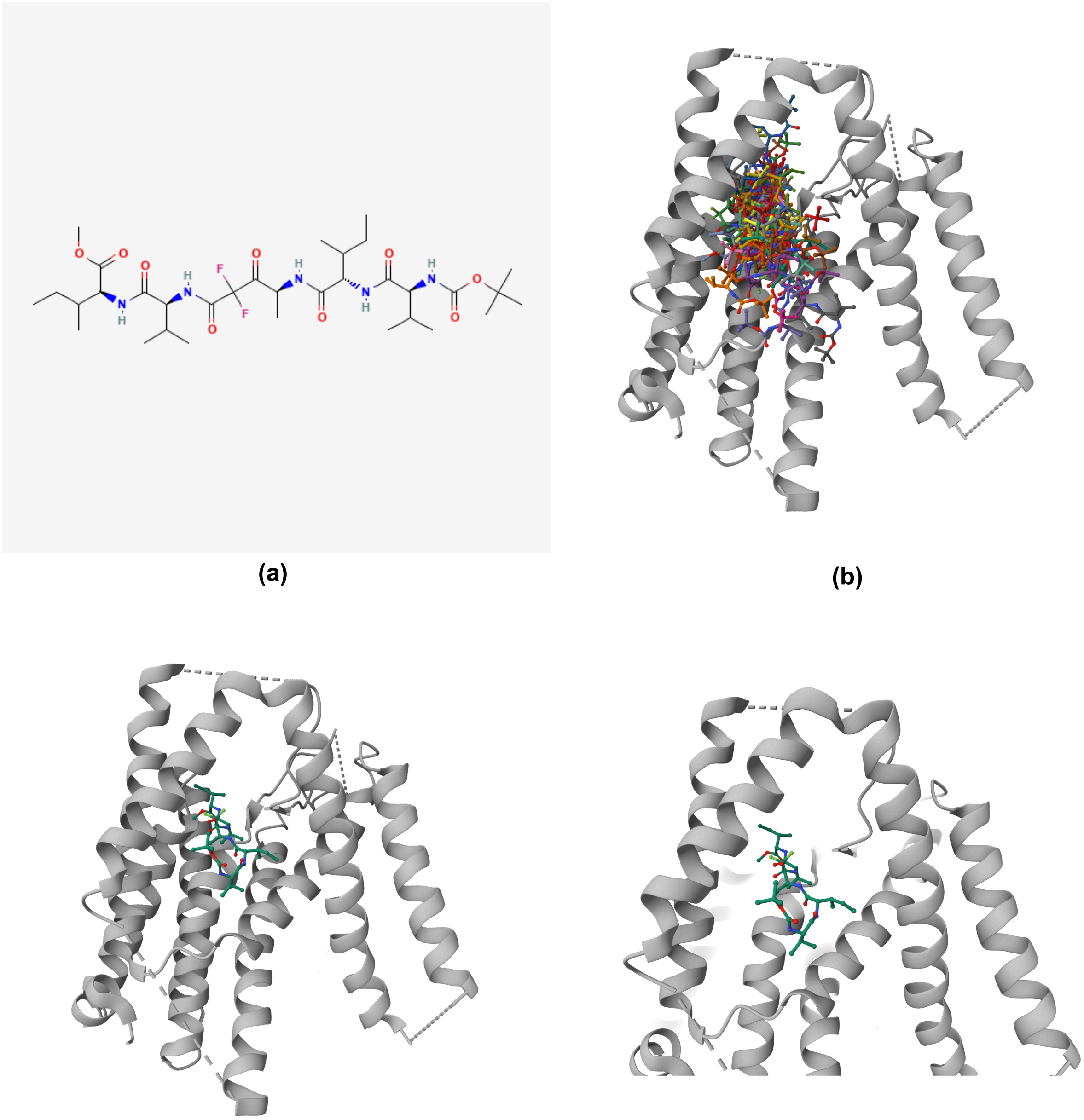


(c) (d)

**Figure S5. Docking analysis of DAPT with Signal Peptide Peptidase.**
(a) Molecular structure of DAPT. (b) Multiple predicted docking conformations generated using DiffDock. (c) Highest-scoring docking pose of DAPT within the predicted binding pocket of Signal Peptide Peptidase. (d) Close-up view of the best docking configuration highlighting ligand positioning within the active site.

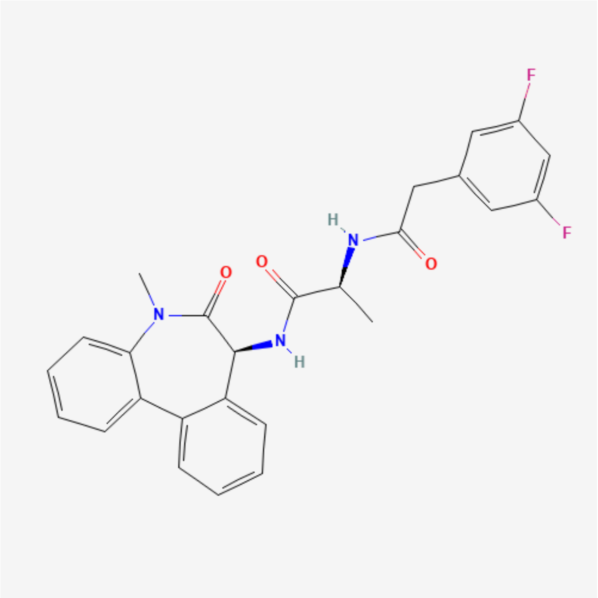

**(a)**

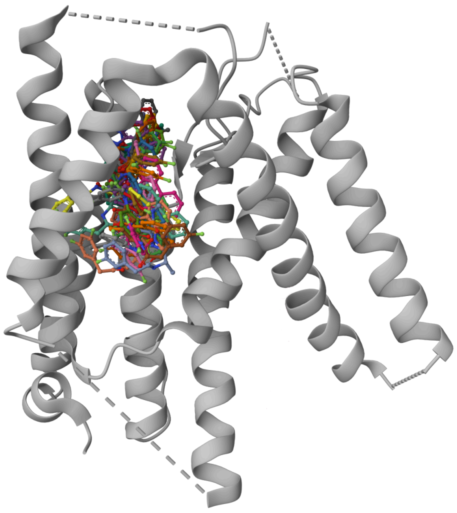

**(b)**

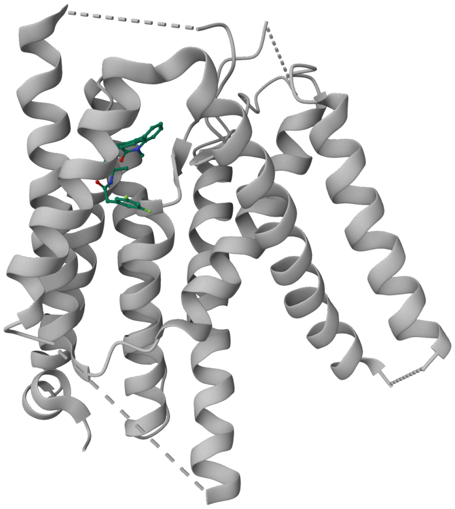

**(c)**

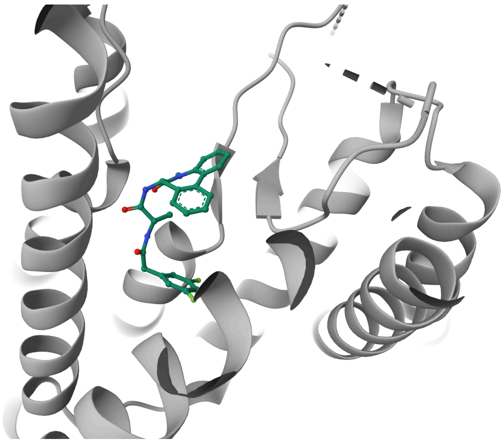

**(d)**

**Figure S6. Docking analysis of N-[(1S)-2-[[(7S)-6,7-dihydro-5-methyl-6-oxo-5H-dibenz[b,d]azepin-7-yl]amino]-1-methyl-2-oxoethyl]-3,5-difluorobenzeneacetamide with Signal Peptide Peptidase.**

(a) Molecular structure of the molecule. (b) Multiple predicted docking conformations generated using DiffDock. (c) Highest-scoring docking pose of the molecule within the predicted binding pocket of Signal Peptide Peptidase. (d) Close-up view of the best docking configuration highlighting ligand positioning within the active site.

#### Performance summary

**Best performing docking candidates.** In Table S2, we present confidence levels provided by DiffDock for all best-docked 6 potential inhibitors. Overall, based on the confidence levels, Compound E shows the most promising binding characteristics for SPP inhibition (-0.28), while among the SP inhibitors, (Z-LL)2-ketone demonstrates better binding potential than L685,458.

**Table S2. Best performing docking candidates for the six potential inhibitor compounds identified for Signal Peptidase (SP) and Signal Peptide Peptidase (SPP). Confidence levels are provided by DiffDock.**

| Compound Name | Target | PubChem CID | Confidence Level ↑ |
|---|---|---|---|
| (Z-LL)2-ketone | SP | 44449123 | -2.05 |
| L685,458 | SP | 5479543 | -2.30 |
| Compound E | SPP | 11306390 | -0.28 |
| DAPT | SPP | 5311272 | -1.11 |
| GSI II | SPP | 44305139 | -2.03 |
| N-[(1S)-2-[[(7S)-6,7-dihydro-5-methyl-6-oxo-5H-dibenz[b,d]azepin-7-yl]amino]-1-methyl-2-oxoethyl]-3,5-difluorobenzeneacetamide | SPP | 11454028 | -0.99 |

### References

(Corso et al., 2023) Corso, Gabriele, et al. "DiffDock: Diffusion Steps, Twists, and Turns for Molecular Docking." International Conference on Learning Representations (ICLR 2023). 2023.
